# Photocycle characterization of a blue-orange cyanobacteriochrome from *Synechococcus* sp. PCC 7002

**DOI:** 10.1101/2022.09.18.508392

**Authors:** Aleksandar Mihnev, Liam Forbes, J. Douglas McKenzie, Richard J. Cogdell, Anna Amtmann

**Author notes:** **Corresponding author: Aleksandar Mihnev E-mail:**. Department of Systems & Computational Biology, Albert Einstein College of Medicine, Bronx, NY 10461, US.

## Abstract

Cyanobacteria employ photoreceptors called cyanobacteriochromes (CBCRs) to sense the colour and intensity of light. The information extracted from the solar spectrum is used for adaptive responses such as optimizing photosynthesis, phototaxis and cell aggregation. cGMP-phosphodiesterase/adenlylate cyclase/FhlA (GAF) domains are the principal light sensors in cyanobacteriochromes. They contain a conjugated bilin chromophore and boast an impressive spectral diversity. Characterizing the spectral characteristics of GAF domains in model strains, such as *Synechococcus* sp. PCC 7002, can open new avenues for optogenetics and biotechnology. Based on sequence analysis we predicted several different GAF domains in this strain. The *SynPCC7002_a0852* gene encodes a single GAF domain with two cysteine residues: one in the conserved α3 helix and one in the conserved DXCF motif. Spectral analysis of recombinant *SynPCC7002_A0852* with phycocyanobilin (PCB) showed that the protein cycles between two states, Po and Pb, which absorb orange and blue light, respectively. Measurements of kinetics identified Po as the dark state of the protein. Acid-denaturation analysis suggested that the 15E isomer of PCB is bound in the (dark) Po state, whereas 15Z is bound the (photoproduct) Pb state. Site-directed mutagenesis and iodoacetamide treatments showed that Cys73 in the DXCF motif is essential for the conversion from Po to Pb. Future experiments dark-purified protein/chromophore versions are required to establish the sequence of events in the photocycle. In summary, *SynPCC7002_A0852* enables orange/blue colour perception in *Synechococcus* sp. PCC 7002 as other CBRCs of this protein family but might contain the energetically higher chromophore isoform in its dark state. Such photocycle has previously been found in ‘bathy’ bacteriophytochromes but not in CBCRs.

## INTRODUCTION

Cyanobacteriochromes (CBCRs) are a family of highly tunable photoreceptors that serve as sensors of light wavelength and intensity in cyanobacteria (Rockwell et al., 2012c). In their simplest form, CBCRs consist of a single *cGMP-phosphodiesterase/adenlylate cyclase/FhlA* (GAF) domain (Rockwell et al., 2015). However, most putative CBCRs are multidomain proteins consisting of several GAFs and other sensory/effector domains, such as the Per-Arnt-Sim (PAS) and histidine kinase (HisK). While several GAF domains and signaling pathways have been characterized (Bezy and Kehoe, 2010; Bhaya, 2016; Hirose et al., 2010, 2013a; Kehoe and Gutu, 2006) the full range of photo-perception mechanisms and signaling targets in cyanobacteria remain to be explored.

The GAF domain is a small globular structure of about 19 kDa. GAF domains can be evolutionarily adapted to sense a variety of environmental parameters (Aravind and Ponting, 1997; Ho et al., 2000; Schultz, 2009), which explains their great ubiquity in nature. CBCR GAF domains usually contain a covalently attached phycocyanobilin (PCB) molecule (Burgie and Vierstra, 2014; Ikeuchi and Ishizuka, 2008; Rockwell et al., 2011, 2013a). The natural spectral properties of the bilin are further tuned by the protein environment and/or by isomerisation (Ishizuka *et al*., 2011; Rockwell *et al*., 2011, 2012a; Chen *et al*., 2012; Rockwell et al., 2013; Yuu Hirose *et al*., 2013; Fushimi *et al*., 2020). The result is a myriad of unique spectral sensitivities: CBCR GAF domains can perceive wavelengths in the UV, visible and far-red regions (Fushimi and Narikawa, 2019). Both CBCRs and phytochromes make use of the GAF domain (Rockwell and Lagarias, 2010). These domains cycle between two spectral states, P_1_ and P_2_, where 1 and 2 indicate the perceived colour. In doing so, CBCR GAF domains tend to measure the ratios of two different colors of light, and unlike phytochromes, they are not restricted to red/far red-absorbing states (Pr and Pfr). The transition between the two holoprotein states is founded upon rotation of the fourth chromophore ring (Chen et al., 2012; Rockwell et al., 2013b; van Thor et al., 2006). Depending on the extent of this rotation around C15, the chromophore itself cycles between a 15Z and a 15E isomer, each being associated with one of the holoprotein P states. In solution, the 15Z state is the more stable of the two (Fushimi et al., 2016). In all currently known CBCRs, the 15Z state of the chromophore seems to be associated with the dark/ground state of the holoprotein. Traditionally, CBCR GAF domains are grouped into families based on sequence similarity. These families are named according to the most prevalent set of colours perceived by their members (e.g. “red-green family”). As the number of spectrally characterized GAF domains grows it becomes evident that these colours rarely apply to every single member. To make a distinction between homology-based annotation and confirmed colour perceptionwe propose the term “chromotype” to denote the latter. The chromotype is defined by the specific combination of spectral colours sensed by a GAF domain, relative to the 15Z and 15E chromophore state, respectively, making no assumptions as to which state is the dark state. For example, a GAF domain with a Pr (15Z) and Pg (15E) configuration will have a Red/Green (R/G) chromotype, and a Pb (15Z) and Po (15E) GAF domain will have a Blue/Orange (B/O) chromotype.

There is evolutionary advantage for organisms to be able to measure the ratios of spectral colours in the environment. These ratios result in a mixed population of P_1_ and P_2_ states, depending on their sensitivity and dark reversion kinetics. Cyanobacteria make use of this readout to tell the time, depth in water column, and presence of other (photosynthetic) organisms. Some of the major responses elicited by CBCRs are chromatic acclimation, phototaxis and cell aggregation. Chromatic acclimation is the process of adapting the type and abundance of light-harvesting pigments to make better use of the current light environment (Kehoe and Gutu, 2006). The classic example is the control of ratio of phycocyanin (PC) and phycoerythrin (PE) in *Fremyella diplosiphon* depending on the ratio of green to red light. This process is partially controlled by the RcaE CBCR in a transcriptional manner (Bezy and Kehoe, 2010). Phototaxis is the process of moving towards or away from a specific light stimulus. The example is the phototactic motion of *Synechocystis* sp. PCC 6803 in response to blue light (Yoshihara and Ikeuchi, 2004). In this example the process is at least partially controlled by the PixJ CBCR, possibly by means of dimerization and interaction with the pili on the membrane. Cell aggregation, biofilm formation and buoyancy changes are all under the control of c-di-GMP levels in *Synechocystis* sp. PCC 6803 (Agostoni et al., 2016). The c-di-GMP levels are linked with the activity of a cellulose synthase, which affects biofilm accumulation (Kawano et al., 2011). This process has been tied to blue light (Agostoni et al., 2013), possibly by an unknown CBCR. The c-di-GMP levels are known to be boosted by GGDEF domains (Galperin et al., 2001; Romling et al., 2013) and diminished by EAL domains (Dow et al., 2006), both of which are common in CBCR proteins. Possible roles of CBCRs and other physiological or developmental responses pathways remains to be elucidated.

In addition to the fundamental knowledge gained from studying the physiological roles of CBCR signaling mechanisms, the spectral characterization of CBCRs has potential for applications in photosynthesis-based biotechnology *Synechococcus* sp. PCC 7002 is already a well characterized, transformable metabolic model organism and a potential chassis for biotechnology. Currently, nothing is known about the spectral properties of its CBCRs. Charting the organism’s responses to different combinations of colours could allow the organism’s metabolism to be controlled without chemicaladditives just by using light. This can be done by tapping into existing CBCR pathways or engineering new ones. The first step is identifying putative CBCRs and characterizing them. Here we describe the identification and characterization of a novel single-GAF domain encoding gene from *Synechococcus* sp. PCC 7002, SYNPCC7002_A0852g, which in the interest of readability we will refer to as Syn0852g henceforth, showing that the protein is a Blue-Orange GAF domain, both in chromotype and sequence similarity, but with an unusual photocycle where the 15E state is the dark state and 15Z is the photoproduct.

## RESULTS

### Recombinant Syn0852 is a blue and orange sensor

A BLAST-P search was performed using the GAF domain amino acid sequence from the CBCR PixJ from *Themosynechococcus elongatus*. The results listed 16 putative proteins from the genome of *Synechococus* sp. PCC 7002 containing at least one GAF domain (Figure 1). Only three GAF domains from these putative proteins exhibited high sequence similarity to known CBCRs, as opposed to GAF domains per se: SynPCC7002_A0689 is likely a member of the well-studied Red-Green (RG) family, SynPCC7002_A0852 is likely a member of the underrepresented Blue-Orange (BO) family and SynPCC7002_A0900 is a predicted member of the CikA (CA) family. Both CA and RG GAF domains seem to be linked with HisK or HAMP-MCP effector domains. The CA family members are currently known to only absorb blue light and have been linked with the cell clock (Schmitz et al., 2000). The RG family is the most spectrally diverse. BO GAF domain photocycles have been characterized in-depth before (Sato et al. 2019; Rockwell et al. 2012a), all of which currently have the B/O chromotype and occur as single domain proteins. Here, SynPCC7002_A0852 (UniProt ID: P32039) was selected for experimental characterization because it seemed like the only candidate from a sequence-and spectrally-conserved family, yet likely to have two distinct spectral P states. This is of interest as any deviation in its spectral properties would be easier to pinpoint based on the sequence. For readability we will refer to SynPCC7002_A0852 as Syn0852g in the following text.

**FIGURE 1:**
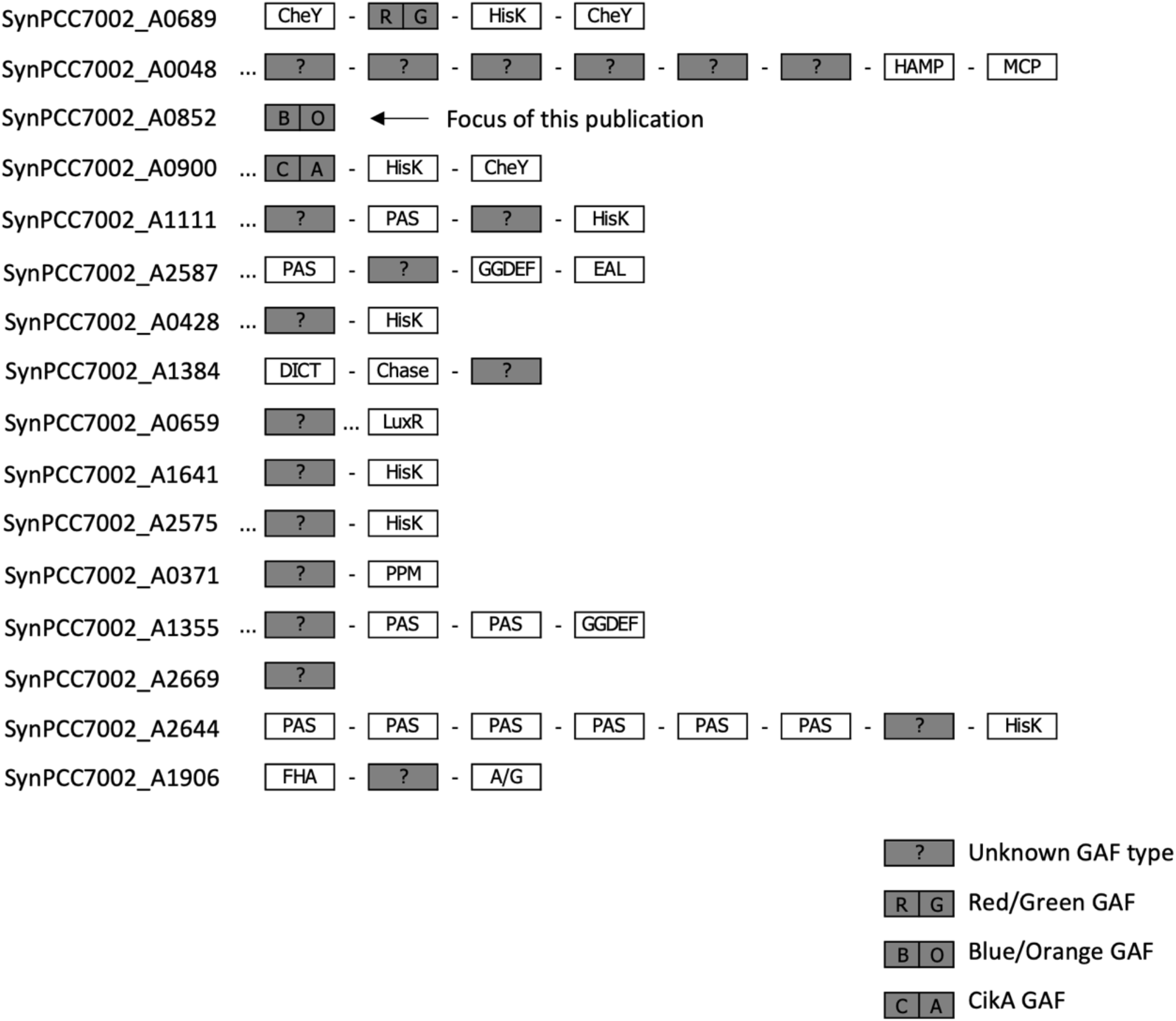
Three CBCR GAF domains are predicted in *Synechococcus* sp. PCC 7002. A BLAST-P analysis was performed against the genome of *Synechococcus* sp. PCC 7002 using the TePixJg sequence as template. For each of the genes listed, the predicted domain structure of the encoded protein is shown from N-terminus (left) to C terminus (right). GAF domains are marked as dark grey rectangles. The predicted GAF domain family, if known, is marked with two letters: RG, BO, and CA. Regions with unrecognized domain signature are marked as “…”. The naming of other domains follows InterPro convention. A/G: Adenylyl cyclase class-3/4/guanylyl cyclase domain. dCach: dCache_1 domain. HOMO: Homodimerization domain. MHY: MHYT domain. S/T-K: Serine/Threonine Kinase domain.

To assess the spectral properties of Syn0852 recombinant protein with integrated PCB was generated in PCB-biosynthesis competent *E. coli* and purified. Initial light treatments were used to monitor photocycling in Syn0852 and the first observations indicated that the protein was reversibly photochromic (Figure 2a). The recombinant protein eluted from the Ni^2+^ resin as a deep blue solution at room temperature. The colour changed from blue to green when exposed to a high intensity halogen bulb through a yellow filter (for filter specifications see Suppl. Figure 2). The green solution seemed stable under ambient light (a combination of natural daylight and fluorescent room light, see Methods) and room temperature (22 °C). Reversion to the blue solution was achieved by replacing the yellow filter with a blue filter (Suppl. Figure 2) or by subjecting the protein to darkness at 4 °C overnight. Absorbance spectroscopy was employed to further investigate the different protein states (Figure 2b,c). The results indicate that the blue solution has a maximal absorbance (λ_max_) around 600 nm, which corresponds to monochromatic yellow/orange (orange hereafter) (Figure 2 d,e). The green solution has a λ_max_ at 440 nm which falls within the blue spectral region (Figure 2 d,e). In the following, the two states are referred to as Po (orange absorbing state) and Pb (blue absorbing state), respectively.

**FIGURE 2:**
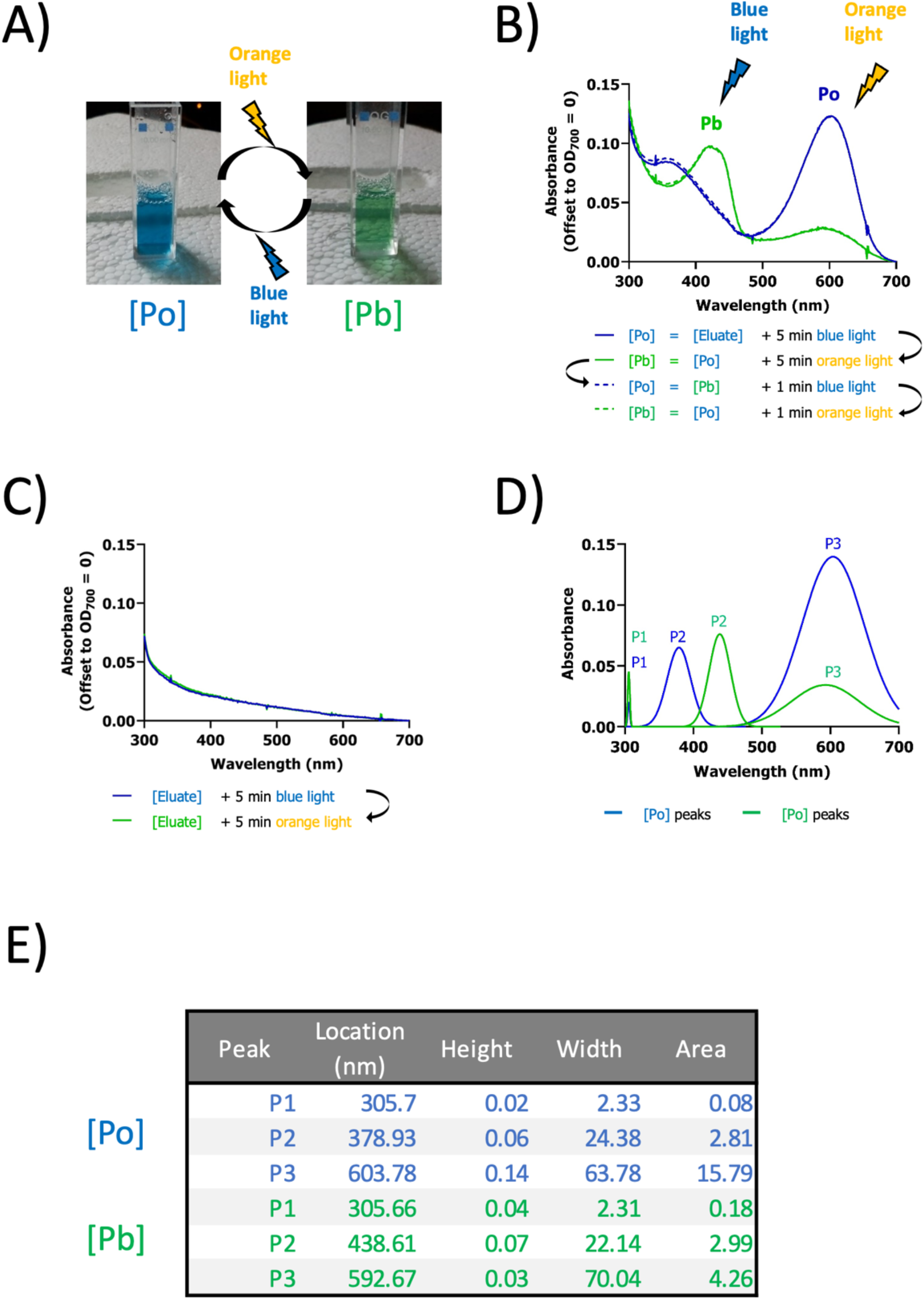
Recombinant Syn0852 with PCB absorbs orange or blue spectral light. Absorbance states of recombinant Syn0852. **A:** A blue solution (Po) was the default state under standard light conditions (see Methods), converting to a green solution (Pb) upon exposure to orange light. Absorbance spectra were recorded after illuminating the protein solution with a halogen-tungsten lamp through colour filters removing wavelengths in the blue spectrum (‘orange’ light) or removing longer wavelengths (‘blue’ light). Emission spectra of the light treatments are detailed in Supplemental Fig. 2. **B:** Absorbance spectra of Syn0852 co-expressed with the chromophore phycocyanobilin (PCB). The sample was illuminated for 5 minutes with blue light (blue line). The same sample was immediately illuminated with orange light (green line). The last sample was also illuminated with the blue light again but for the shorter time of 1 min (dashed lines) and with a orange light again for 1 min. The protein states producing the absorption spectra shown were called Po (blue lines) and Pb (green lines). **C:** Absorbance spectra of Syn0852 without PCB. **D:** Composite peaks (P1-3) predicted from the spectra shown in B). Po peaks are shown in blue. Pb peaks are shown in green. **E:** Composite peak data from D).

### The Po state is the state of the protein

We observed that Syn0852 always purified as a deep blue solution (Po state). The Pb state also appeared stable under standard light conditions with little to no noticeable reversion. After leaving samples in either the Po state or Pb state over night in the dark at room temperature or in the refrigerator, we always found them in the Po state the following morning. Therefore, we hypothesized that the Po state serves as the dark/ground state of the protein. To confirm and quantify this observation, we employed two approaches. The first approach was steady state spectroscopy with quasi-continuous illumination. We used pulses of light immediately followed by measurements in an iterative manner (Figure 3a,b). Both transitions showed systematic changes in A600 in the timescale of 1 min, consistent with cumulative light exposure, without obvious evidence of additional relaxation processes. The second more traditionally kinetic approach utilized a single-wavelength dark-reversion kinetics at A420 nm (Figure 3c-e). In this method, we synchronized the protein population to the Po state via a long exposure to blue light, followed by a varying pulse of yellow light to motivate the protein to the Pb state. We then kept the protein in the spectrometer for the duration of the study whilst monitoring A420. We tested three different timescales: 16 seconds (Figure 3c), 16 minutes (Figure 3d), and 7 hours (Figure 3e). While the first two timescales provided no reliable insights due to low R2 scores, the dark-reversion on the scale of 7 hours (Figure 3e) showed a clear shift from the Pb state towards the Po state, with a high R2 score of 0.992 and a rate constant (k) of -1.56 x 10-4 for the dark reversion. This strongly suggests a dark decay in the Pb -> Po direction, consistent with preliminary observations.

**FIGURE 3:**
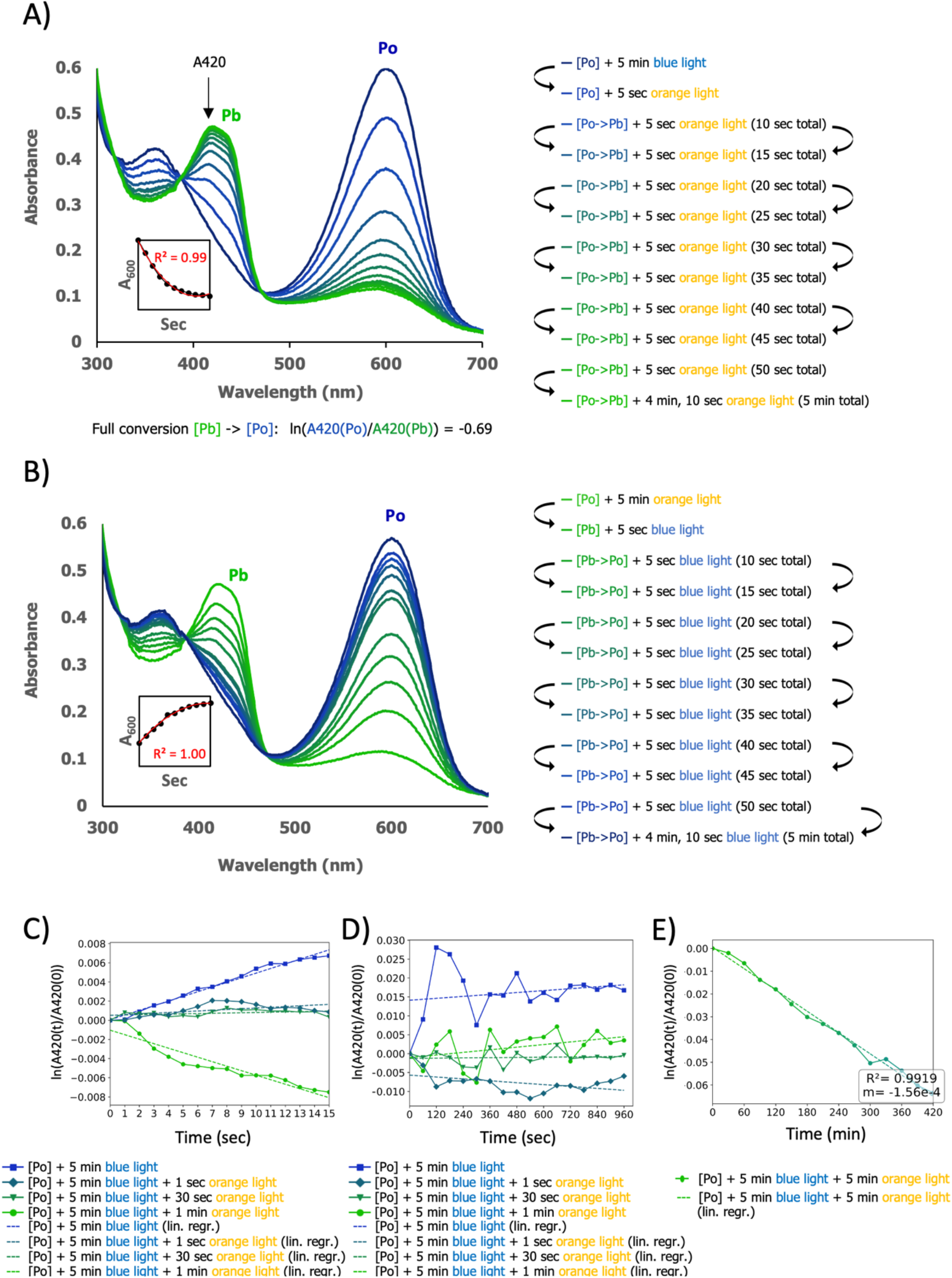
Syn0852 time-based relaxation. Po/Pb conversion timescales and kinetics. All experiments were performed at room temperature (22°C). **A:** Conversion Po -> Pb. Protein eluate in the Po form was exposed to blue light for full conversion to Po. Sequential orange light exposures determined saturation duration to Pb via uv/vis spectroscopy. **B:** Conversion Pb -> Po. Protein eluate in the Po form was exposed to orange light for full conversion to Pb. Sequential blue light exposures determined saturation duration to Po via uv/vis spectroscopy. **A, B:** Insets shows ýA600 (≤60 sec) with a fitted polynomial of the second order. **C:** Dark reversion kinetics at short timescales. Full Po conversion was ensured by blue light exposure. Various durations of orange light (0s, 1s, 30s, 60s) motivated the protein toward the Pb state. A420 readout every 10th sec, averaged every 1 sec. R2 values: 0.98, 0.31, 0.16, 0.90. k values (ΔY sec⁻¹): 4.86×10⁻⁴, 7.69×10⁻⁵, 2.88×10⁻⁵, -4.70×10⁻⁴. **D:** Dark reversion kinetics at medium timescales. Data from B) shown at longer timescales. R2 values: 0.04, 0.20, 0.01, 0.16. k values (ΔY sec⁻¹): 4.25×10⁻⁶, -4.18×10⁻⁶, 5.04×10⁻⁷, 5.76×10⁻⁶. **E:** Dark reversion kinetics at longer timescales. Full Po conversion followed by 5 min orange light (●). R2 = 0.99. k value (ΔY sec⁻¹): -1.56×10⁻⁴ (ΔY min⁻¹). Readout via A420, averaged every 30 min.

### The dark (Po) state contains 15E PCB

Acid-denaturation spectroscopy was used to interrogate the chromophore species and configurations in the Po and Pb states (Figure 4). The assay involves denaturing the protein allowing direct spectral observation of the chromophore and assigning C15 double-bond configurations (15Z/15E). Under blue light the absorbance spectrum of the denatured protein was reminiscent of the native protein in Po with two absorbance peaks at 358 and 580 nm. Subsequent exposure of the denatured Po sample to white light shifted the orange light absorption from 580 nm to red at 666 nm. This observation is indicative of PCB relaxation from 15E to 15Z as inferred by acid-denaturation assays from other CBCRs (Fushimi et al., 2016). Under orange light the protein is in the Pb state. The denatured Pb protein showed the same absorbance spectrum as the denatured Po sample following white light illumination. The spectrum of the denatured Pb sample did not change upon subsequent exposure to white light. These results indicate that the Po state contains phycocyanobilin as 15E isomer (15E PCB), whereas the Pb state contains 15Z PCB. The data from the deconvoluted peaks is shown in Suppl. Figure 3. The chromotype of Syn0852, expressed as the 15Z/15E colour combination, is therefore blue/orange (B/O). This is in accord with other CBCRs from the same gene family. However, Syn0852 is unusual insofar as the Po (15E) state is the dark state of the holoprotein, meaning that the protein will eventually fall back to this state in the absence of light stimulation. In solution, the 15E isomer of PCB eventually decays into the 15Z isomer (van Thor et al., 2006) and accordingly the dark states of all CBCRs characterized to date, including other members of the BO family, contain 15Z PCB. The novel result for Syn0852 suggests the presence of one or more stable bonds between of Syn0852 and the chromophore in the 15E state, making it the energetically-preferred state.

**FIGURE 4:**
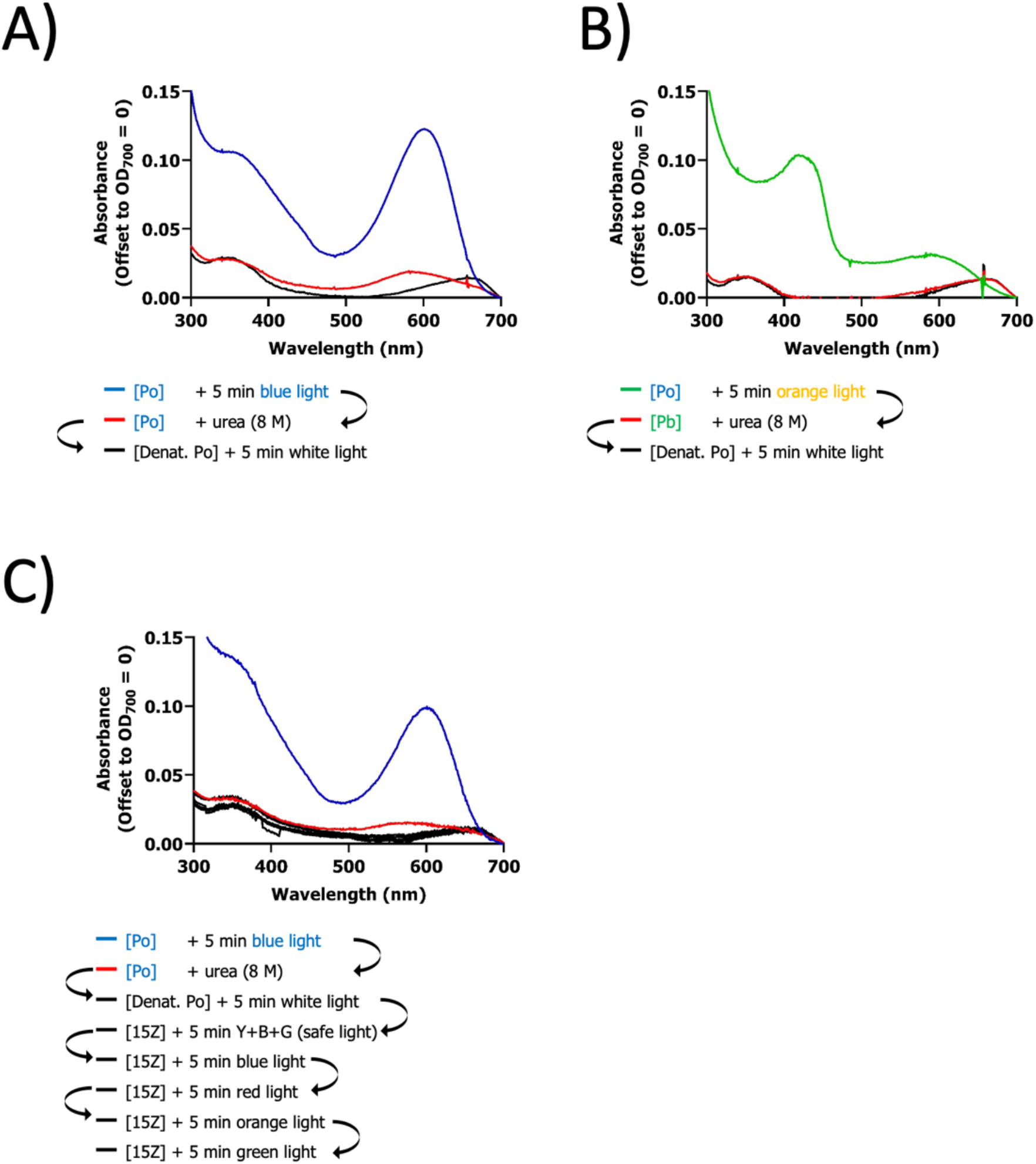
The Po/Pb states of Syn0852 correspond to the 15E/15Z isomers of phycocyanobilin. Recombinant Syn0852 was expressed in the presence of PCB and purified as a blue solution under standard light conditions (see Methods). Aliquots of the solution were exposed for 5 minutes to ‘blue’ or ‘orange’ light to convert all protein to Po (blue line) or Pb (green line), respectively. A, B: Absorbance spectra of Syn0852 in Po state (A) or Pb state (B) before (blue/green lines) and after (red line) addition of urea at a final concentration 8 M. A third absorbance spectrum (black line) was measured after illuminating the urea-treated samples for 5 minutes with white light. C: As in A with additional light treatments applied after white light illumination of the denatured protein (grey lines).

### Cysteine 73 is required for the photocycle

Blue light absorption in photoreceptors is known to be achieved by means of a Cys residue interacting with the conjugated system of the chromophore, forming a thiol bridge to C10 of the bilin (Rockwell, 2015). Syn0852 contains only two cysteine residues. One, Cys101, is in a conserved position within helix α3. This residue is generally associated with constitutively binding to PCB at ring A (Ikeuchi and Ishizuka, 2008). The second residue, Cys73, is located within the DXCF motif and is suspected to yield the blue light sensitivity of the Pb state. Recombinant C73S and C73A protein was generated as before and both proteins eluted as purple solutions. Absorbance spectroscopy of the mutant proteins (Figure 5) indicated an abolished photocycle, two photostates with the same spectral characteristics, or the protein being stuck in a new or intermediate state. The data for the deconvoluted peaks are shown in Suppl. Figure 4. The results indicate that Cys73 is essential for tuning the spectral properties and/or the transition between photostates. The Syn0852 chromotype is B/O (15Z/15E) which corresponds to that of other members of the Blue-Orange family. Cys73 does not explain any differences in the dark state isoform because all proteins of the Blue-Orange family have this conserved residue.

**FIGURE 5:**
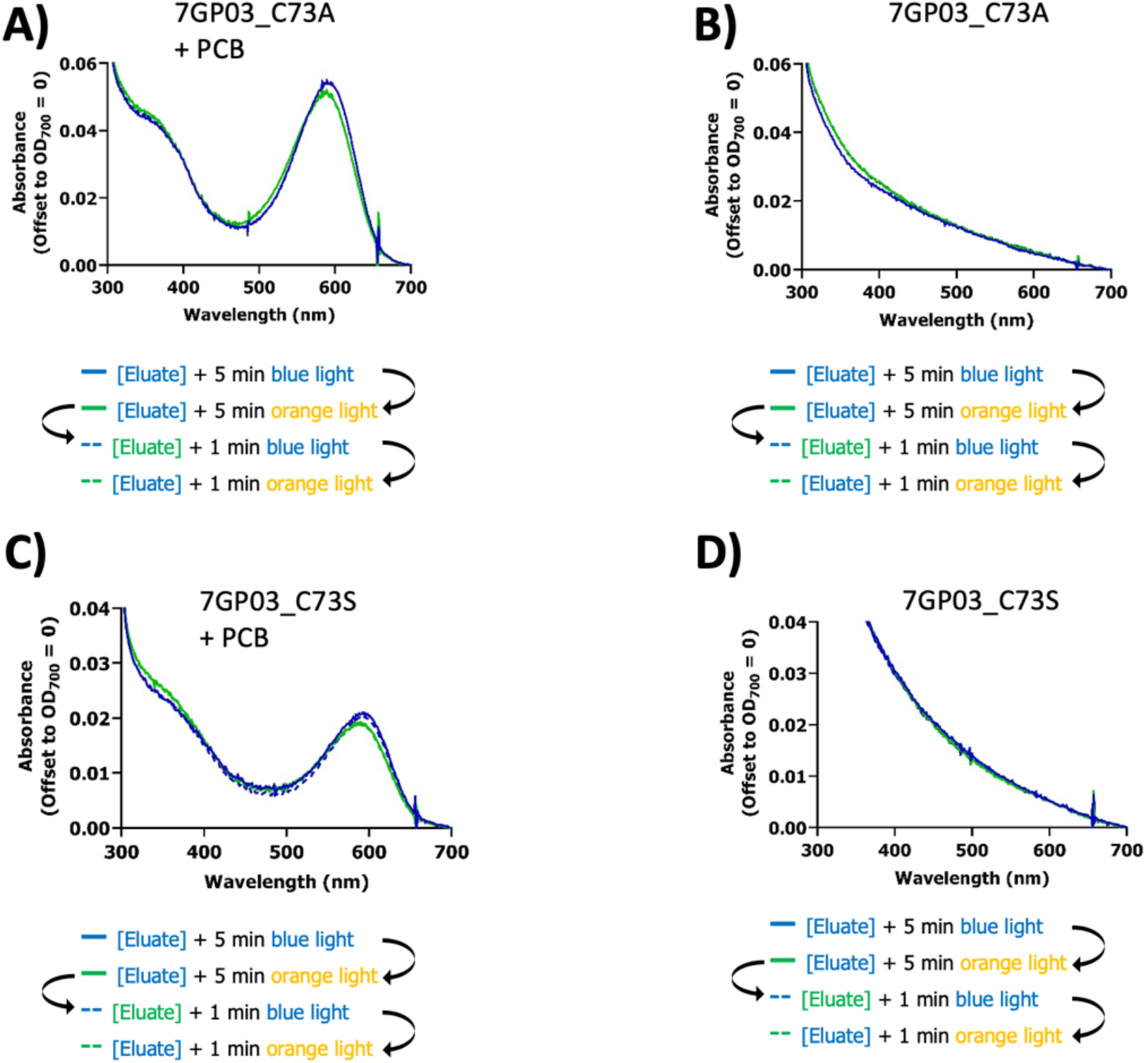
Cys73 is essential for the photocycle. Absorbance spectra of mutant versions of Syn0852 in which cysteine 73 was replaced with alanine (Syn0852_C73A) or serine (Syn0852_C73S) by site-directed mutagenesis. The recombinant proteins were expressed in the presence or absence of PCB and purified under standard light conditions (see Methods). Absorbance spectroscopy was carried out after treating the protein with ‘blue’ light (blue lines) or ‘orange’ light (green lines) for 1 minute (dashed blue and dashed green lines) or 5 minutes (blue and green lines). A, B: Absorbance spectra of Syn0852_C73A with PCB co-expressed (A) or not (B). C, D: Absorbance spectra of Syn0852_C73S with PCB co-expressed (C) or not (D).

### Cys73-C10 link is not essential for 15E (Po) -> 15Z (Pb) transition

B/O CBCRs contain the DXCF motif. Traditionally, the 15Z/15E interconversion is thought to be the main and first photocycle event in DXCF CBCRs, followed by the formation of a second Cys linkage at C10 of PCB (Hardman et al., 2020). To better understand the photocycle order of events in Syn0852 we applied acid-denaturation spectroscopy to the mutant proteins Syn0852_C73S and Syn0852_C73A (Figure 6). The results from the peak deconvolution are shown in Suppl. Figure 5. The inability of the mutants to form the Pb state (shown in Figures 4,5) is now accompanied by the denatured protein shifting its absorption from orange to red upon white light treatment. The results of this experiment indicate that PCB remains as the 15E isomer in both mutant proteins, meaning that removal of Cys73 abolishes the ability of the chromophore to isomerize. Moreover, the experiment suggests that the C73A and C73S mutant versions of the protein were not in the 15Z/Pb state. It is likely they were arrested in a changed 15E/”Po” state and/or an intermediate state. Thus, while these observations confirm that Cys73-mediated activity is essential for the photocycle they are not sufficient to prove that formation of the Cys73-C10 link is the first event of the photocycle. In future, mutant proteins would need to be expressed and purified in complete darkness to avoid intermediate state.

**FIGURE 6:**
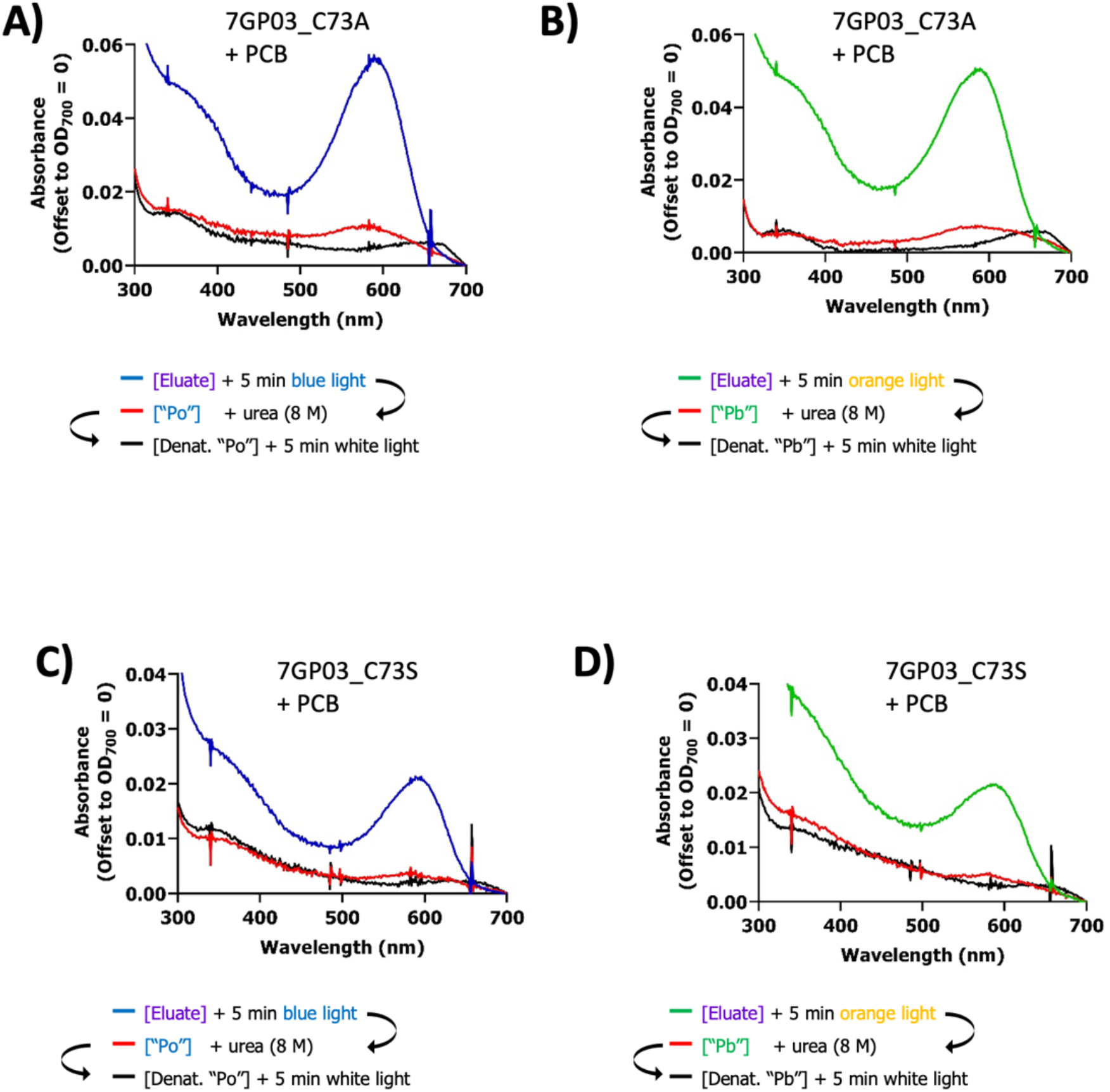
The role of Cys 73 in the photocycle is essential for 15E-15Z isomerization. Absorbance spectra of denatured mutant versions of Syn0852 replacing cysteine in position 73 with alanine (Syn0852_C73A) or serine (Syn0852_C73S) were generated by site-directed mutagenesis, the recombinant proteins were expressed in the presence or absence of PCB and purified under standard light conditions (see Methods). Aliquots of the solution were exposed for 5 minutes to ‘blue’ or ‘orange’ light to try and convert all protein to Po (blue line) or Pb (green line), respectively. A, B: Absorbance spectra of Syn0852_C73A in “Po” state (A) or “Pb” state (B) before (blue/green lines) and after (red line) addition of urea at a final concentration 8 M. A third absorbance spectrum (black line) was measured after illuminating the urea-treated samples for 5 minutes with white light. C, D: Absorbance spectra of Syn0852_C73S in “Po” state (A) or “Pb” state (B) before (blue/green lines) and after (red line) addition of urea at a final concentration 8 M. A third absorbance spectrum (black line) was measured after illuminating the urea-treated samples for 5 minutes with white light.

The Cys73-PCB link is the primary suspect for a Cys73-mediated activity in the photocycle of the wild-type CBCRs (Rockwell et al., 2012b). To interrogate this hypothesis and address the pitfall of the previous experiment we used a pharmacological approach. Iodoacetamide (IAM) alkylates any free Cys residues and therefore prevents formation of thiol bridges with other residues/molecules (Gurd, 1972). This has been used with CBCRs before (Enomoto et al., 2012; Narikawa et al., 2014). Synchronizing the protein population to Po was done by a prolonged exposure to blue light first. Absorbance spectroscopy then showed that treatment with IAM impaired the ability of the proteins to transition from Po to Pb (Figure 7). The deconvoluted peak data from this experiment is shown in Suppl. Figure 6. These results showed that that a Cys73-PCB link was essential for transitioning into Pb.

**FIGURE 7:**
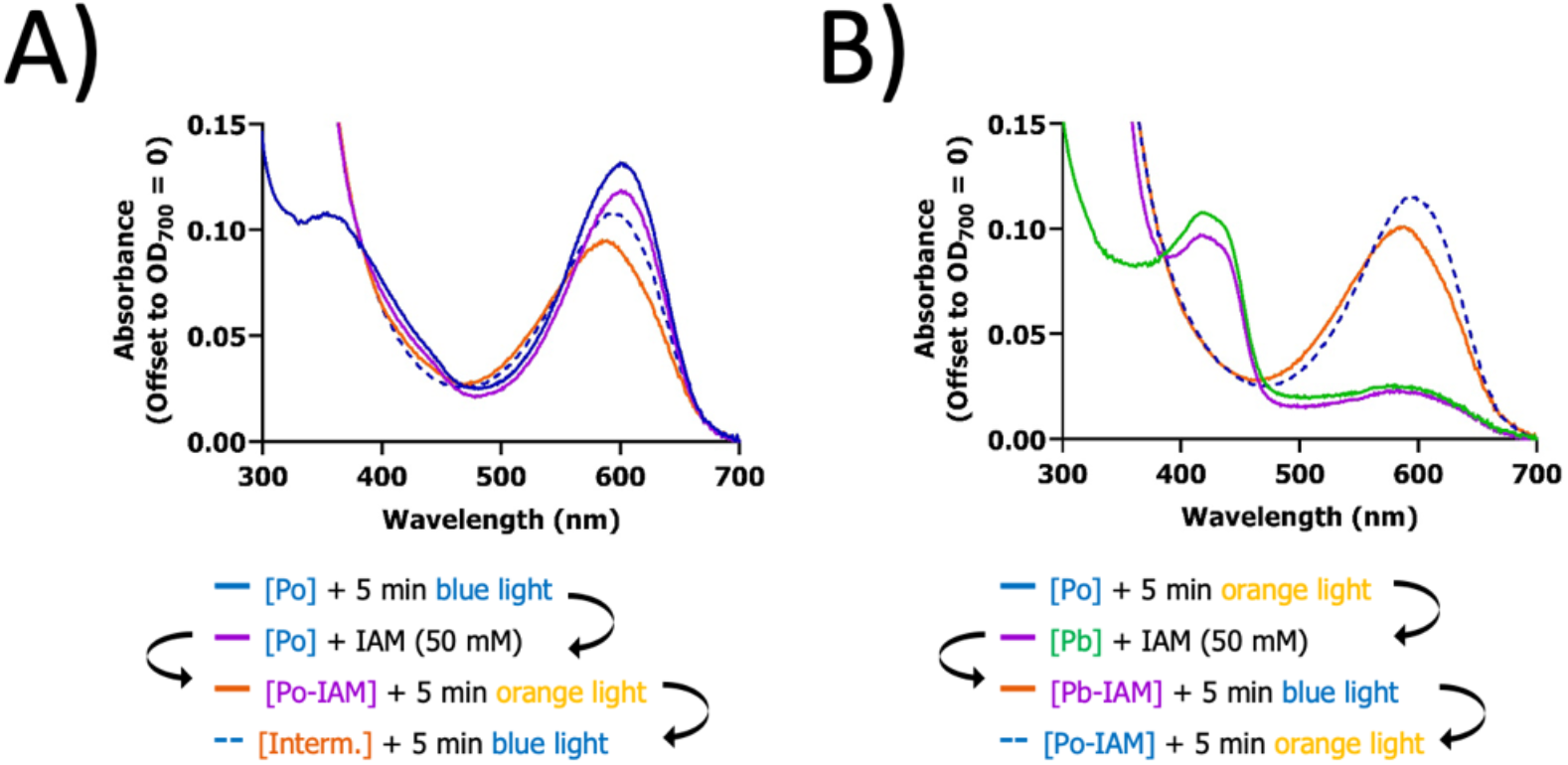
A thiol bridge is likely formed between Cys73 and C10 of PCB in the Po to Pb transition. Absorbance spectra of Syn0852 after treatment with iodoacetamide (IAM). Recombinant Syn0852 was expressed in the presence of PCB and purified as a blue solution under standard light conditions (see Methods). Aliquots of the solution were exposed for 5 minutes to ‘blue’ or ‘orange’ light to convert all protein to Po (A, blue line) or Pb (B, green line), respectively. IAM was then added to a final concentration of 50 mM, immediately followed by an absorbance scan (purple line in A and B. Next, the samples were exposed to 5 min of ‘orange’(A) or ‘blue’ light (B) to drive the protein into the opposite state, and absorbance spectra were measured (orange line in A, dashed blue line in B). Finally, the protein was motivated back to its original state by applying either ‘blue’ (A) or ‘orange’ light (B), and absorbance spectra were measured (dashed blue line in A, orange line in B).

Acid-denaturation spectroscopy was performed on IAM-treated Syn0852 (Figure 8a). The control experiment (Figure 8b) contained the Po form of Syn0852, treated with IAM and subsequently denatured with urea. The peak deconvolution is shown in Suppl. Figure 7. The lack of significant relaxation of 15E PCB to 15Z in the control indicated the presence of the 15E isomer in most of the proteins and ruled out any unexpected effect from IAM. Interestingly, the Po to Pb transition was mostly associated with 15Z PCB in the presence of IAM. In other words, the Cys73-PCB link is not essential for 15E -> 15Z, suggesting that the ring rotation is the first photocycle event (consistent with other CBCRs).

**FIGURE 8:**
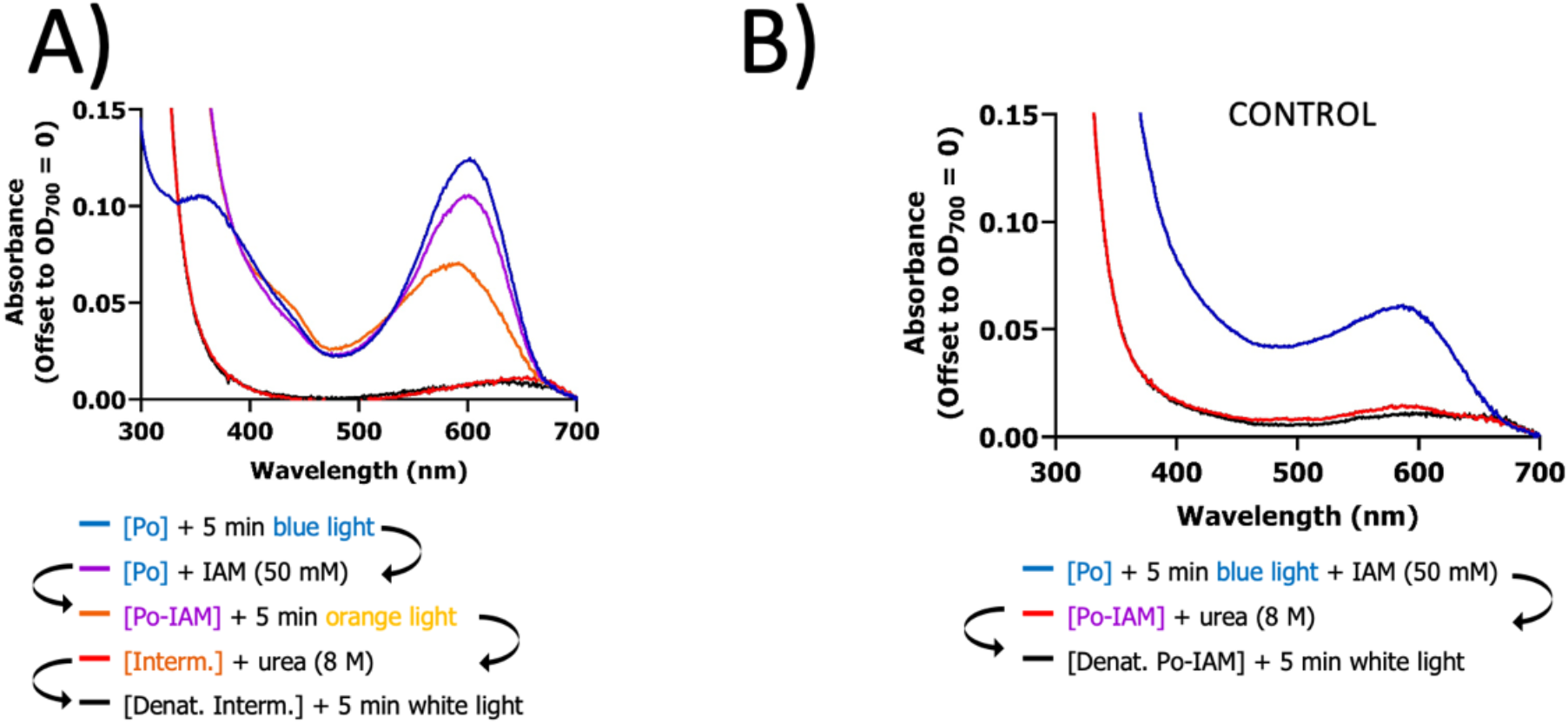
The presence of Cys 73 alone is insufficient to drive the photocycle. Absorbance spectra of denatured IAM-treated Syn0852. Recombinant Syn0852 was expressed in the presence of PCB and purified as a blue solution under standard light conditions (see Methods). An aliquot of the protein was exposed to 5 min ‘blue’ light to take it completely to Po. A: Absorbance spectra of protein before (blue line) and after (purple line) addition of IAM to a final concentration of 50 mM. The IAM-treated sample was then illuminated for 5 min with ‘orange’ light to take it to the intermediate state (orange line). Next, urea was added to a final concentration of 8 M, immediately followed by another absorbance scan (red line). The final absorbance scan was carried out after illuminating the denatured protein +PCB for 5 min with white light (black line). B: Control absorbance spectra of IAM treated Syn0852 in the Po state (blue line) exposed to urea (red line) and white light (black line) using the same concentration and duration as in A.

In contrast, during the equivalent experiment with Cys73 mutants the chromohphore remained in the 15E state. This indicates that, in addition to forming a link to C10 of PCB, Cys73 is sterically important to the photocycle.

## DISCUSSION

A unique feature of Syn0852, different to all previously characterized CBCRs, including Blue-Orange ones, is that the dark (ground) state of the holoprotein contains PCB as the 15E isomer. In the Blue-Orange family of CBCRs the Po state also contains 15E PCB, just like in Syn0852. However, the Po state has not yet been observed as the dark state of the holoprotein. Site-directed mutagenesis showed that blue light sensitivity of the Pb state is linked with the activity of Cys73 in the DXCF motif as is the case for other CBCRs with similar wavelength sensitivity and the same motif. Additional pharmacological treatments and mutagenesis studies indicated that thiol bridge formation between Cys73-C10 of PCB in the Pb state does not explain why the dark state (Po) is the more energetically favorable despite containing 15E PCB and no thiol bridge. The combined results imply that additional residues and/or bonds stabilize this isomer in Po of Syn0852. Although this result is new in the context of CBCRs, such opposing photobiological properties have been seen in bacteriophytochromes (Karniol & Vierstra, 2003).

Our experiments prove that Syn0852 is a B/O CBCR where the bilin is bound as 15E and 15Z in the Po and Pb states, respectively. We also provided evidence that the Po state is the dark state of the holoprotein via first-hand observations and a kinetic study. Despite of our proof that Cys73 is needed for the Pb state using mutagenesis and pharmacological studies, we could not confidently establish a link between this residue and the energy landscape of the protein. Moreover, we could not establish the order of events in the photocycle, relative to Cys73 activity. In order to address this, the mutant protein would need to be purified and studied in complete darkness.

We can theorize as to the likely suspects behind the Po state being the dark state by comparing Syn0852 to other members of the BO family, sharing high sequence similarity but not exhibiting such behaviour. The relative heights and positions of the absorbance spectra of Syn0852 (439 nm, 604 nm) are almost identical to those of NpF4973g (434 nm, 602 nm). NpF4973g from *Nostoc punctiforme* is a member of the BO family thus also having a Po and Pb state (Rockwell et al, 2012a). Similarly, the Po state also contains 15E PCB and the Pb state contains 15Z PCB with a thiol link. In the case of NpF4973g the Pb state is the dark state of the holoprotein. The two proteins have over 80% amino acid sequence similarity. Most of the difference is due to the N-terminal extension in NpF4973g, which is unlikely to participate in the photocycle. If this region is excluded from the analysis, the sequence similarity is 92%. The remaining 8% difference probably accounts for the opposing energy landscape of the Po states. Fushimi *et al*. (2020) identified six key residues in the binding pocket of GAF domains with an effect on the chromotype. The positions of these key residues can be mapped to all currently characterized CBCR GAF domains. Syn0852 is unique among all currently characterized GAF domains in having an aspartate residue (Asp 114) in one of these six key positions (Asn 109 equivalent in NpF4973g and Asn 109 TePixJg). The net negative charge of the aspartic acid under physiological conditions may create a charge attraction with the fourth ring of PCB, thus stabilizing the 15E isomer in Po. Additional site-directed mutagenesis in both proteins should be carried out in the future to test this hypothesis.

Further interrogation of the unusual Syn0852 could reveal the basis for its unusual energy landscape and lead to its potential physiological function and signaling pathway. In particular, mutagenesis of other residues within the vicinity of the chromophore will reveal the mechanism stabilizing the 15E chromophore state, making it more energetically favorable than 15E. Other proposed studies will involve substituting the chromophore with different versions of the bilin, which may be supplemented with NMR and/or X-Ray crystallography. Finally, taking other BO CBCRs and replacing key residues (for example, position 114 to Asp) in their binding pockets to resemble Syn0852, followed by spectroscopic analysis could be used to further elucidate the mechanism.

## Supporting information

Data for figures

Script for kinetics figures. Run as .py

## ACKNOWLEDGEMENT

We offer special thanks to Prof. David Kehoe (Indiana University Bloomington) for our initial introduction to the field of cyanobacteriochromes. We extend our gratitude to Prof. Kai Hong Zhao (Huazhong Agricultural University) for the pACYC-ho1-pcyA plasmid and to Prof. John Christie (University of Glasgow) for consultation and use of equipment in his laboratory.

We also thank the following people, in order of their last names, for specialist training provided: Dr Ross Eaglesfield (University of Glasgow) for protein expression, Dr Mary Ann Madsen (University of Glasgow) for cyanobacteria culturing and handling, Dr Sakharam Waghmare (University of Glasgow) for FPLC training.

This work was supported by the Industrial Biotechnology Innovation Centre (IBioIC, PhD studentship to AM) and the Biotechnology and Biological Sciences Research Council (BBSRC; B/V509279/1 PhD studentship to LF, BB/N018508/1 and BB/09825/1 to AA).

## AUTHORS’ CONTRIBUTIONS

AM carried out all experimental work and data analysis, and wrote the draft manuscript, LF contributed to data interpretation and writing, JDM and RJC co-supervised the project, AA led the project and co-wrote the manuscript.

## METHODS

### Strains and plasmids

*Synechococcus* sp. PCC 7002 was obtained from the Pasteur Culture Collection (PCC). It was kept as DMSO-cryopreserved stocks used as inoculum for working cultures. The pET21c-GPR-EGFP vector was provided by Dr Ross Eaglesfield (University of Glasgow). The pACYC-ho1-pcyA vector was a gift from Prof. Kai Hong Zhao (Huazhong Agricultural University). pACYC-ho1-pcyA provides the machinery for *E. coli* cells to produce the phycocyanobilin (PCB) chromophore which self-ligates into CBCRs.

### Molecular cloning

Genomic (gDNA) was extracted from 250 µl of cell pellet of *Synechococcus* sp. PCC 7002 grown in liquid culture (true OD_730_ = 1-3) according was to the method for cyanobacterial gDNA extraction from Tamagnini et al. (1997).

Syn0852 without the endogenous STOP codon was amplified from *Synechococcus* sp. PCC 7002 gDNA by PCR using the Syn0852-FULL-F6/R4 primers and Phusion DNA polymerase (NEB). The pET21c-GPR-EGFP vector was PCR-linearized with pET-F/R and pre-digested with HindIII-HF and XhoI (NEB). The insert was ligated to the vector in a 3:1 ratio (insert:vector) using T4 ligase (NEB). This introduced C-terminal 6xHis tag to Syn0852. The pET-Syn0852-6His vector was transformed into and selected from One Shot Top 10 competent cells (Invitrogen). Plasmid minipreps were done using the QIAprep Spin Miniprep Kit (QUIAGEN).

Site-Dircted Mutagenesis (SDM) was performed by subjecting the pET-Syn0852-6His vector prep to PCR with Syn0852-C73S/A-F/R primers. 1 uL of DpnI (NEB) was added directly to the PCR reactions for 1 hr at 37°C, followed by repeating this step once more. The product was assessed on agarose gel electrophoresis and transformed into TOP10 competent cells as described above, followed by plasmid miniprep with QIAprep Spin Miniprep Kit (QUIAGEN).

Plasmid minipreps were sequenced by first being PCR-amplified using the T7-Prom-F/T7-Term-R primers and cleaned up using the ExoSAP-IT PCR Product Cleanup kit (Thermo Fisher, 78200). Sanger sequencing was then carried out by SourceBioscience for verification.

All commercial materials were used according to the manufacturers’ instructions.

### Expression and purification of recombinant protein

Recombinant Syn0852 protein was made by first co-transforming pET-Syn0852-6His and pACYC-ho1-pcyA into competent BL21 Star DE3 cells (Invitrogen) and selecting transformants with the relevant antibiotics. Two 500 mL cultures of LB medium were then inoculated with the double-transformed cells in 2L conical flasks in the presence of the relevant antibiotic selection. These were grown at 37°C and 180 rpm until OD_600_ = 0.6. The cultures were then cooled to 20°C and induced with 0.5 mM IPTG overnight. The pellets were harvested on the following day and suspended in ice-cold 50 mM Tris-Cl (pH 8.0), 500 mM NaCl, 10-20 mM imidazole, 10% (v/v) glycerol, 0.1%(v/v) Triton X-100 in addition to DNase (0.04 u/mL) and protease inhibitor cocktail. The cells were then lysed on ice using a sonicator and centrifuged at 4 °C and 16 000 x g for 20 min. The supernatant was loaded onto a pre-equilibrated Ni-affinity column. The 3 mL resin was then washed with 200 mL 50 mM Tris-Cl (pH 8.0), 150 mM NaCl, 10-20 mM imidazole, 10% (v/v) Triton X-100. The protein was subsequently eluted with 8 mL 50 mM Tris-Cl (pH 8.0), 150 mM NaCl, 200 mM imidazole, 10% (v/v) glycerol. The selected fractions were pooled together and dialyzed twice at 4oC overnight using a 5000 MW cutoff tubing in 5L 50 mM Tris-Cl (pH 8.0), 150 mM NaCl. The protein was concentrated to 2 mL. Successful generation of recombinant Syn0852 was verified with Zn^2+^ fluorescence and Western blot (Suppl. Figure 1).

### SDS-PAGE, Western blot, Zinc assay

Protein samples were mixed 1:1 (v/v) with 100 mM Tris-Cl (pH 6.8), 4% (w/v) Sodium dodecyl sulphate (SDS), 20% (v/v) glycerol, 3 mg/mL bromophenol blue. The samples were then boiled for 15 min before being subjected to SDS-PAGE using 15% (v/v) acrylamide, 375 mM Tris-Cl (pH 8.8), 0.1% (w/v) SDS, 0.05% (w/v) ammonium persulfate (APS) at 140 V (DC) for 1-2 hours. The bands were visualized by soaking into Coomassie blue, followed by de-staining with dH2O. Gels for Western blotting/Zinc assays were not stained. Instead, the bands were transferred onto a nitrocellulose membrane using the dry blotting method for 1 h at 10 V (DC). The membrane was hen incubated with blocking solution containing 5% (w/v) milk powder in TBST containing 137 mM NaCl, 2.7 mM KCl, 19 mM Tris-Cl (pH 7.4), 0.1% (v/v) Tween 20. After 1 h the membrane was aspirated and a solution of monoclonal α-6xHis antibody (Cell Signalling, 2366) was prepared according to the manufacturer’s instructions, suspended in blocking solution. The antibody was incubated with the membrane at 4°C overnight. The membrane was then washed three times with blocking solution before being incubated with an α-mouse IgG conjugated to horseradish peroxidase (HRP) (Cell Signalling, 7076) in the same way. The membrane was then washed three times with TBST and the bands were visualized using a chemiluminescent kit for HRP (Thermo Scientific, 34075) according to the manufacturer’s instructions. Zinc assays were done according to the method in Kuwasaki *et al*. (2019). This involves submerging the nitrocellulose membranes in a solution of ZnCl_2_ for 30 min, followed by visualization under UV light.

### Ambient light conditions

The ambient light environment consisted of a mixture between warm-white fluorescent lights and sunlight (typically cloudy weather, year-round at 55.8617° N and 4.2583° W). Sample emission spectra taken in the lab are shown in Suppl. Figure 8. The total light intensity at working height was about 25-35 µmol m^2^ s^-1^. Unless stated otherwise in-text, the reader should assume work was done under these conditions, in the presence or absence of additional light stimulation. The light had no effect on the temperature, as the lab was air conditioned. Where an additional high-intensity light source was used (e.g. halogen bulb ± filters in spectroscopy experiments), additional air-circulation was provided in the form of two PC fans, around 15 cm of the sample.

### Spectrometry

Absorbance spectra were taken on a Perkin Elmer Lambda 40 UV/Vis spectrometer. The slit size was set to 0.5 nm. The scanning speed used was 960 nm/min. The UV lamp change was at 326 nm. The resolution was 1 nm. A quartz cuvette was used for all measurements. The protein concentration for most experiments was 0.5 mg/mL. Light treatments were done by adjusting the distance to the halogen lamp (after adding the colour filter) so that the intensity at the level of the sample was always 1000 µmol m^2^ s^-^_1._

### Data analysis

Peak deconstruction was done by importing the spectral data into Igor Pro 8.4 (WaveMetrics) and using the binomial smoothing function (set to 100), followed by the Multipeak Fittin 2 algorithm.

## ACCESSION ID

Syn0852 UniProtKB Accession ID: P32039

---

**SUPPL. FIGURE 1.**
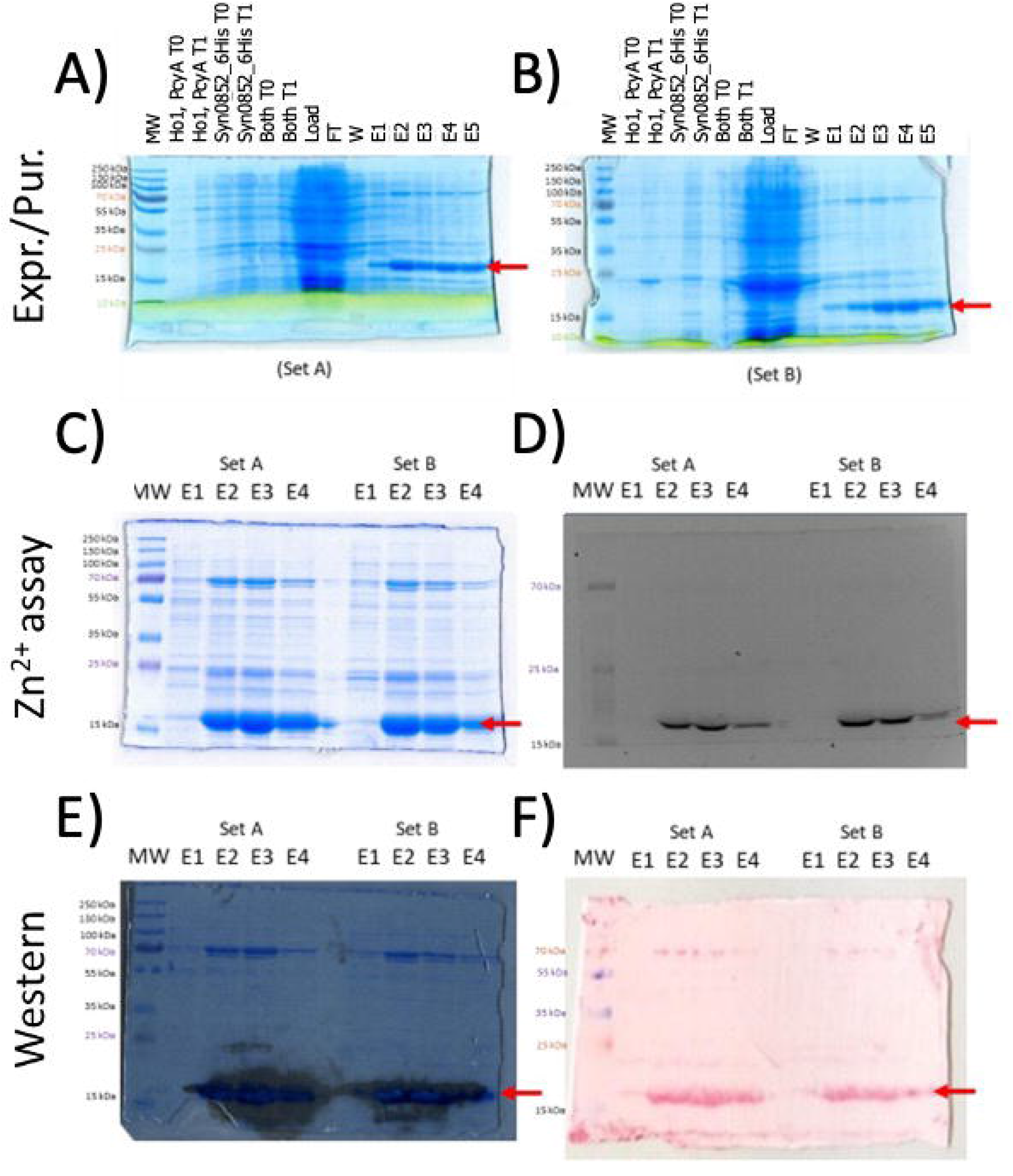

**SUPPL. FIGURE 1.**
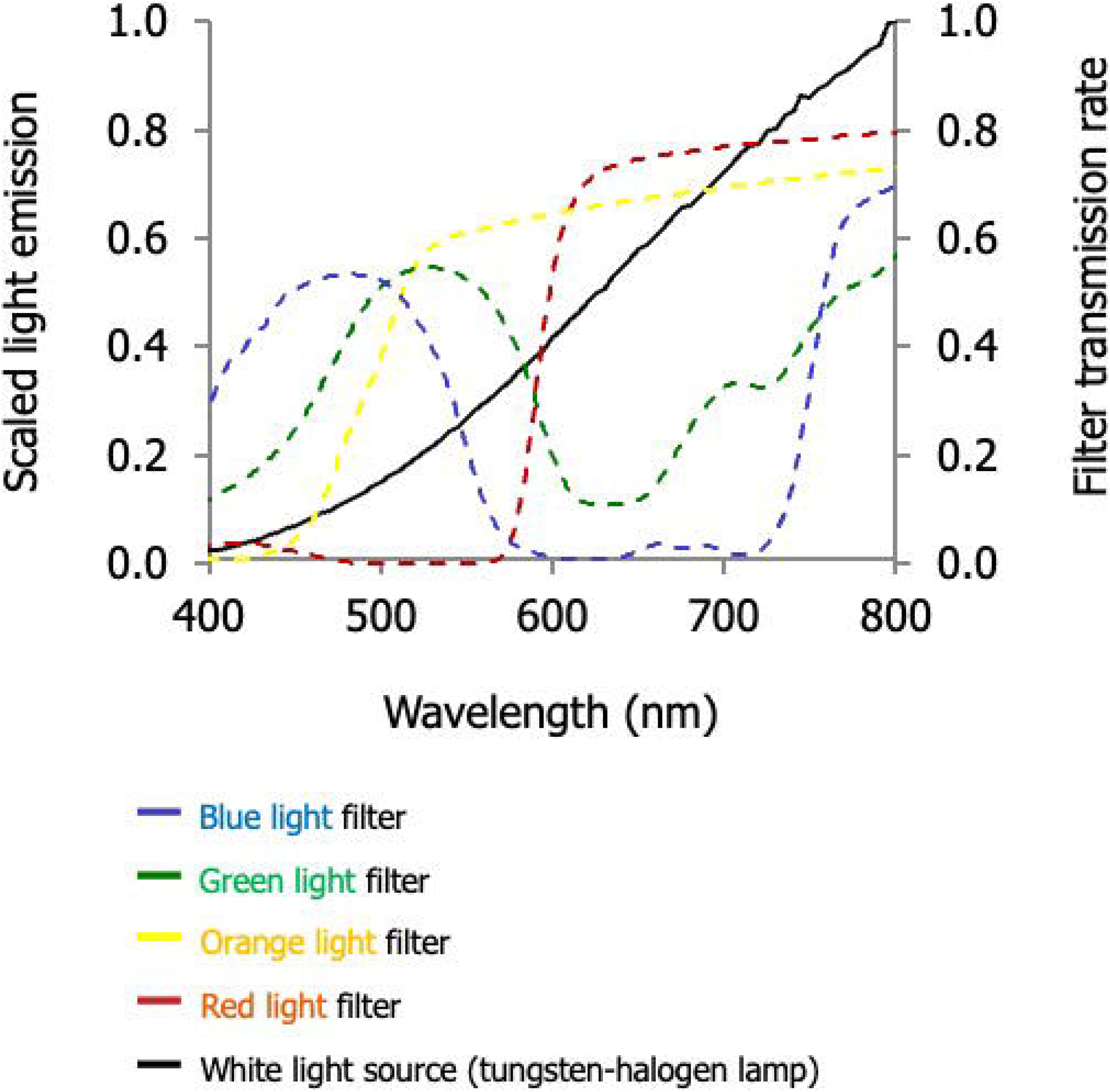

**SUPPL. FIGURE 3.**
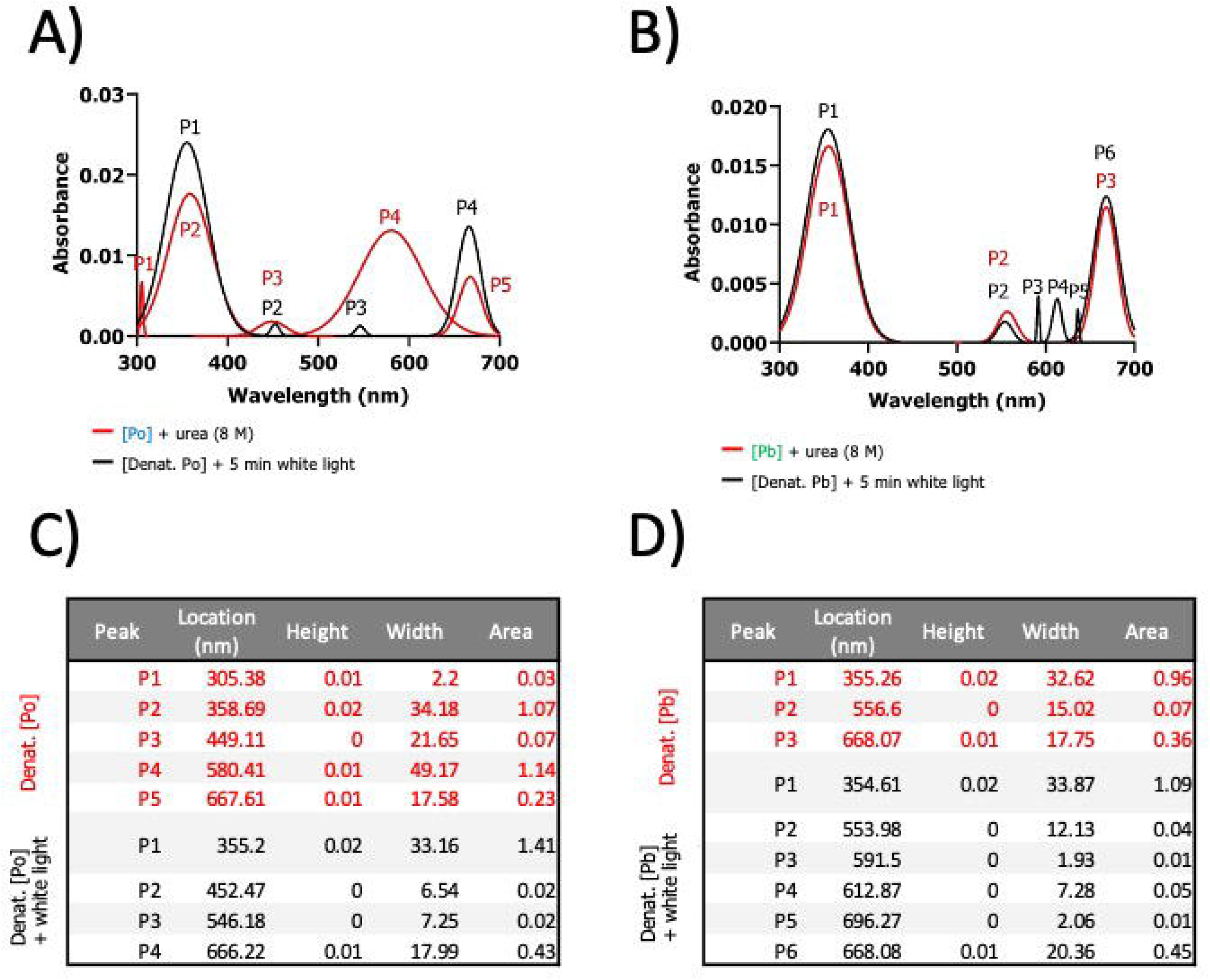

**SUPPL. FIGURE 4.**
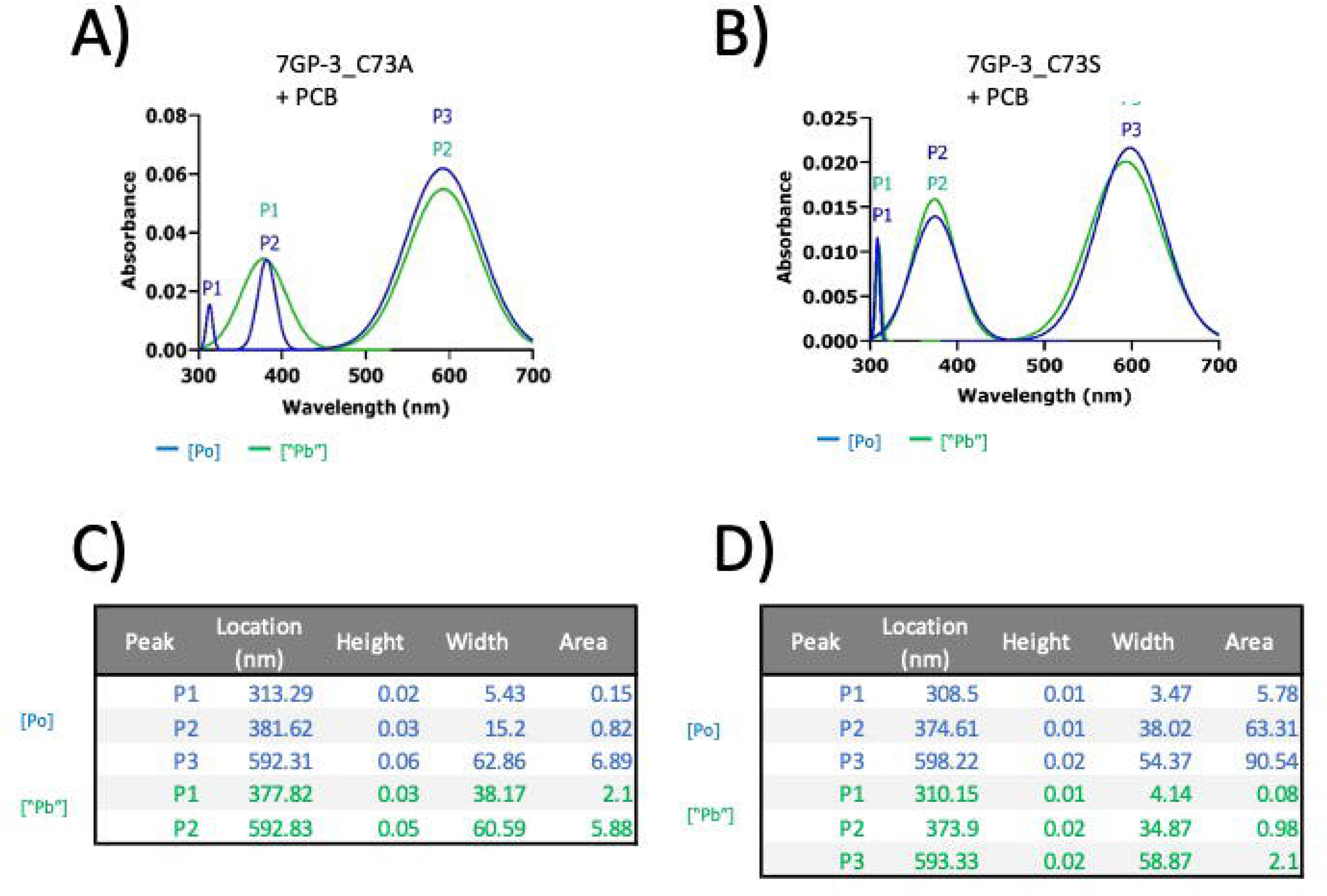

**SUPPL. FIGURE 5.**
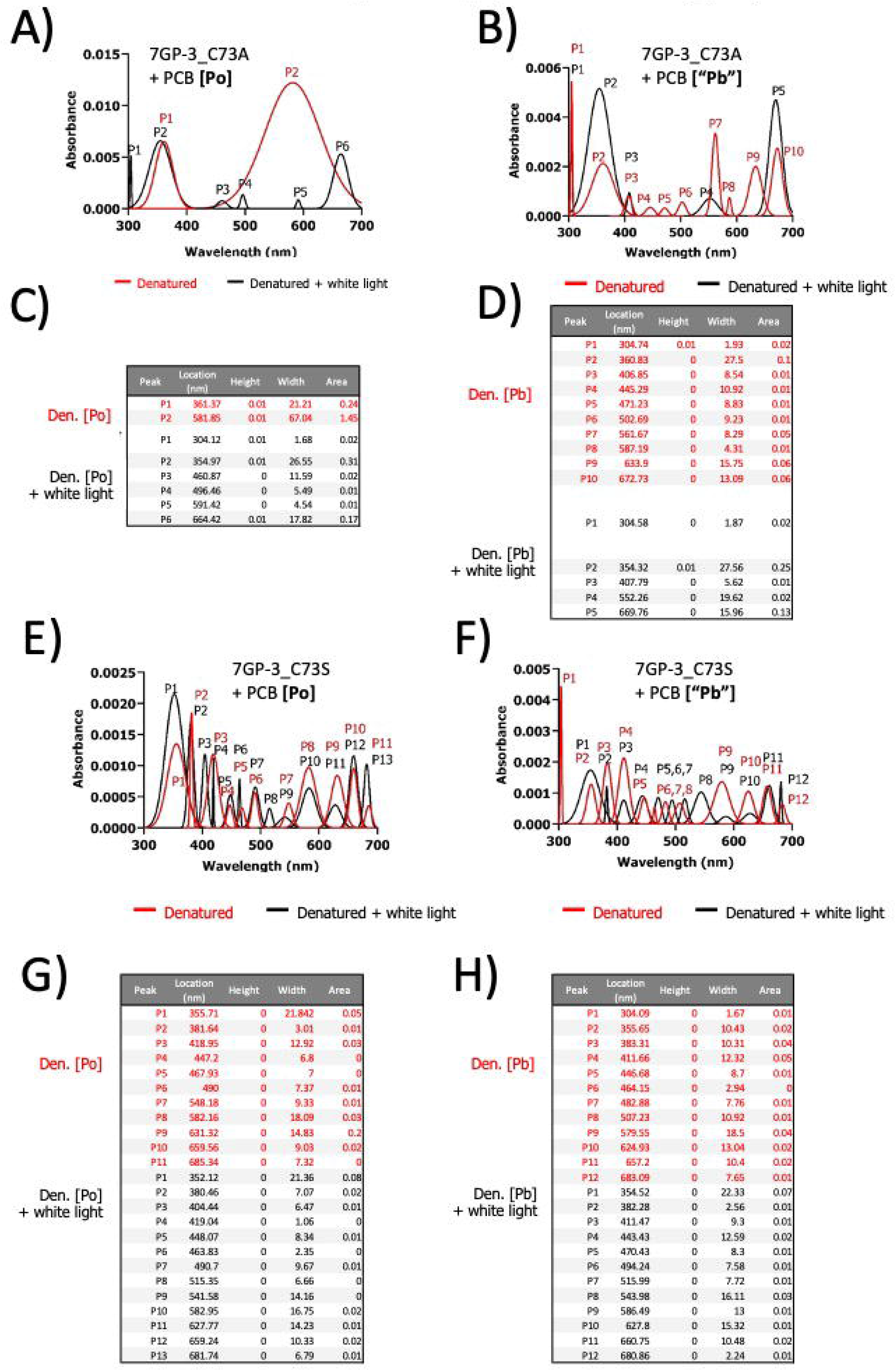

**SUPPL. FIGURE 6.**
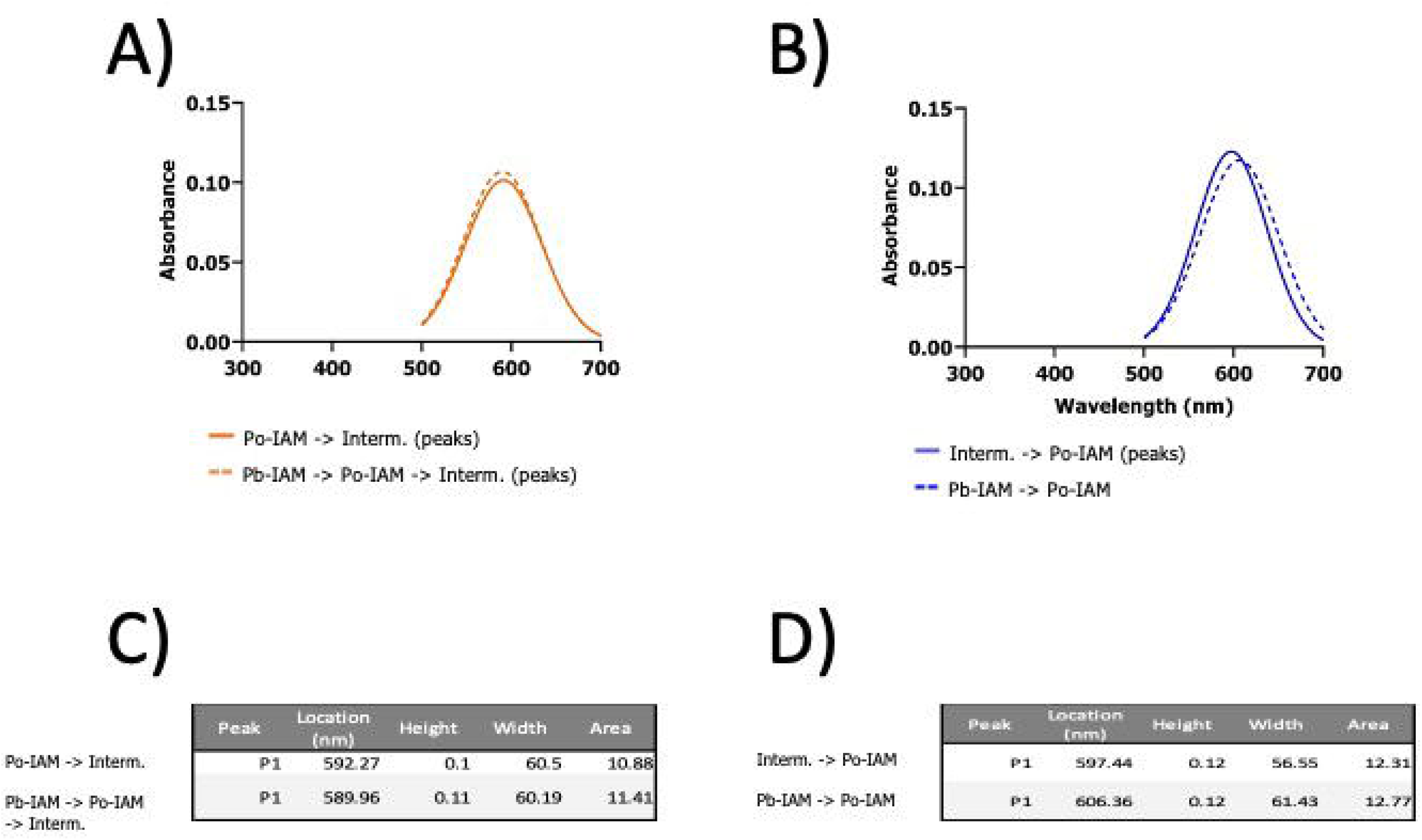

**SUPPL. FIGURE 7.**
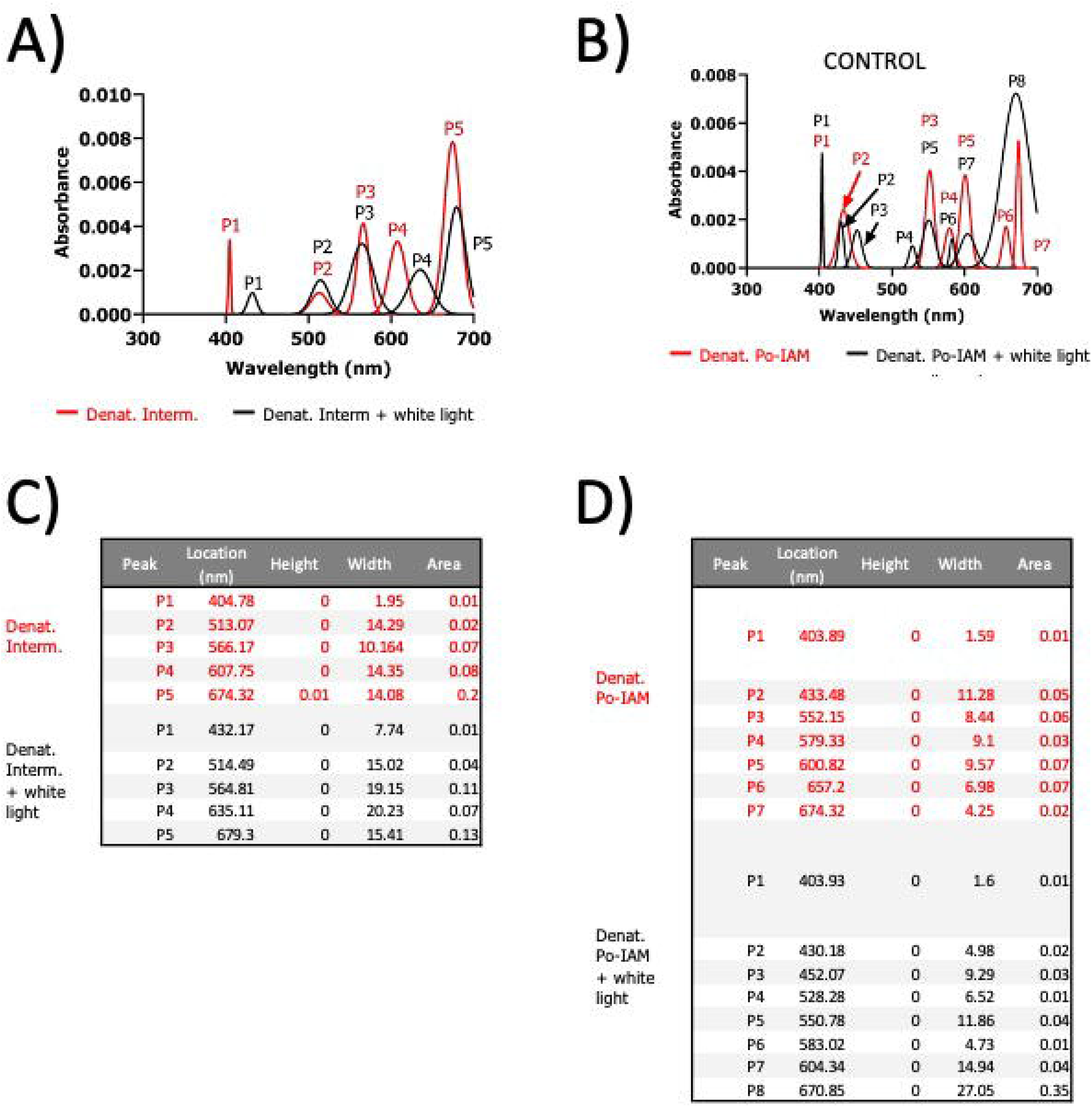

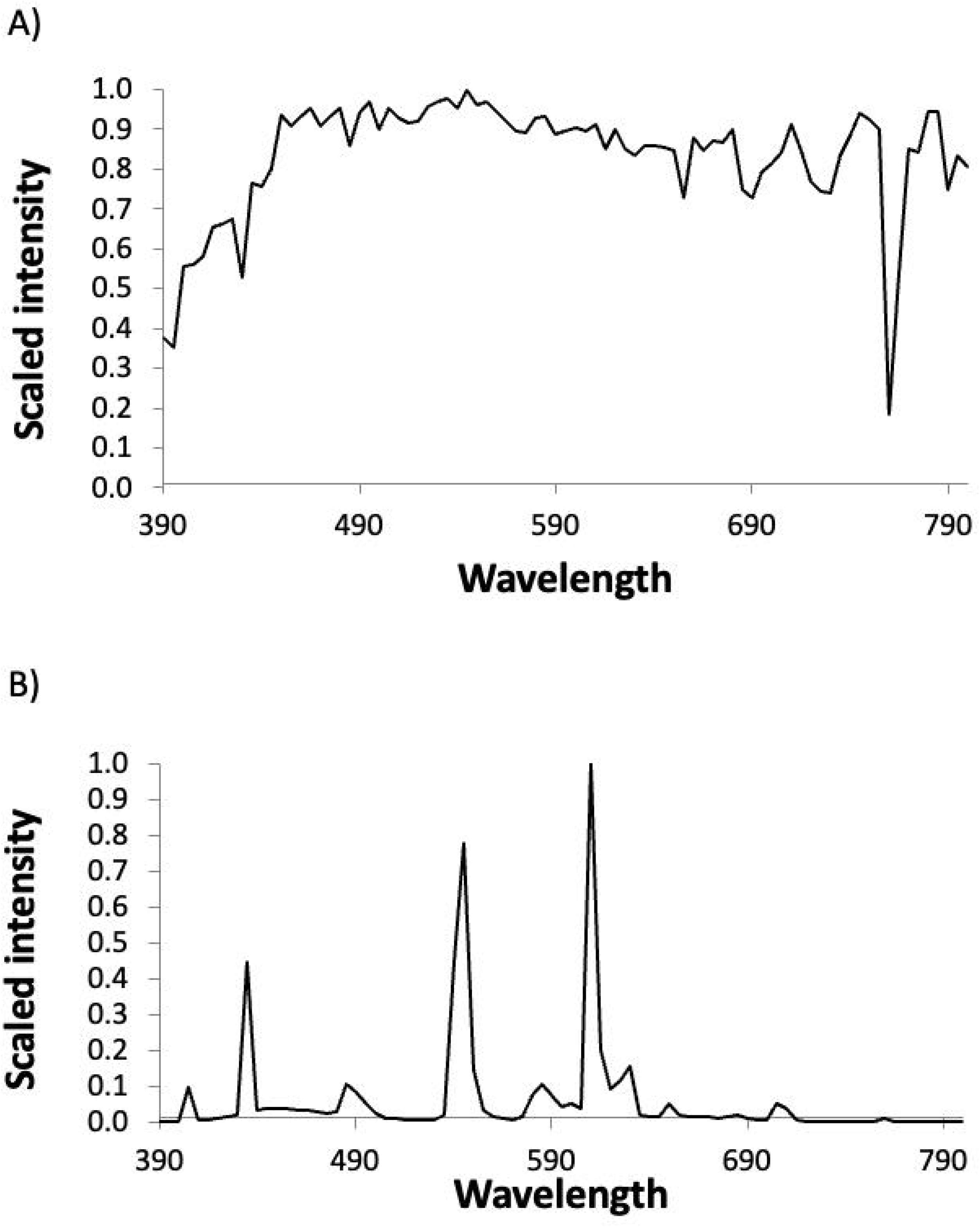

