## Supporting figures for "Photocycle characterization of a blue-orange cyanobacteriochrome from *Synechococcus* sp. PCC 7002"

**SUPPORTING INFORMATION**

**Recombinant protein purification**

SDS-PAGE, Zinc assay and Western blot gels from the purification of 7GP-03 are shown in Supplementary Figure 1.

**Light sources and colour filters**

The emission spectrum of the halogen-tungsten light source and the transmission spectra of the colour filters are shown in Supplementary Figure 2.

**Primer sequences**

| | |
|---|---|
| 7GP-03-FULL-F6: | 5'- GTAT<u>AAGCTT</u>ATGTTTGATCGAAGTCTTAATCGAG-3' |
| 7GP-03-FULL-R4: | 5'- GATC<u>CTCGAG</u>GTCCATGAGTTGTAAATCACC-3' |
| 7GP-03-C73S-F: | 5'-GGGGCCGATGATAGTTTTAATGGTG-3' |
| 7GP-03-C73S-R: | 5'-CACCATTAAAACTATCATCGGCCCC-3' |
| 7GP-03-C73A-F: | 5'-GGGGCCGATGATGCTTTTAATGGTG-3' |
| 7GP-03-C73A-R: | 5'-CACCATTAAAAGCATCATCGGCCCC-3' |
| pET-F: | 5'-GTTCCTCGAGCACCACCACCACC-3' |
| pET-R: | 5'-GGTAAAGCTTTATGTATATCTCCTTCTTAAAGTTAAAC-3' |
| T7-Prom-F: | 5'-TAATACGACTCACTATAGGG-3' |
| T7-Term-R: | 5'-GCTAGTTATTGCTCAGCGG-3' |

**Peak deconstruction**

Peak deconstruction analysis was carried out for the spectroscopy data obtained in this work.

Supplementary Figure 3 shows the deconstructed peaks from Figure 4.

Supplementary Figure 4 shows the deconstructed peaks from Figure 5.

Supplementary Figure 5 shows the deconstructed peaks from Figure 6.

Supplementary Figure 6 shows the deconstructed peaks from Figure 7.

Supplementary Figure 7 shows the deconstructed peaks from Figure 8.

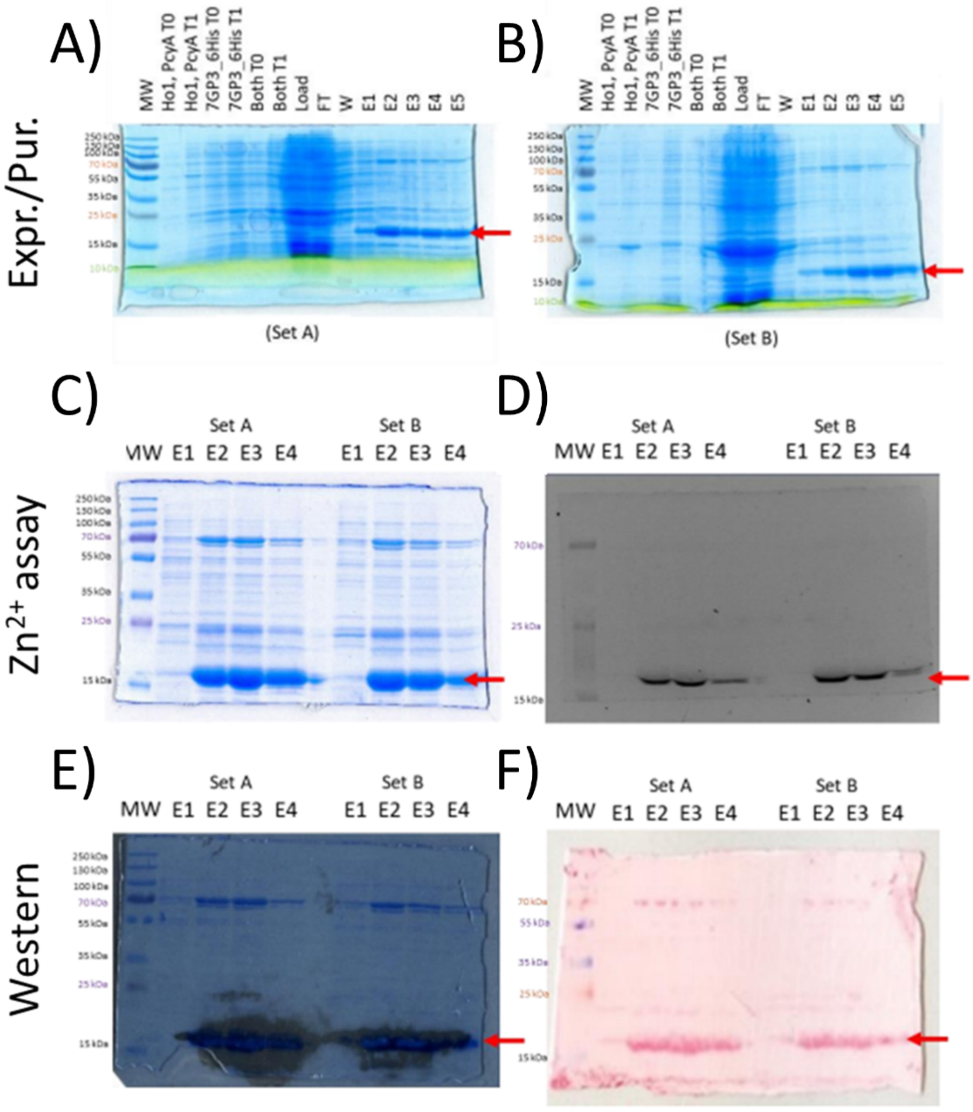


SUPPL. FIG. 1: **Expression and purification of recombinant Synechococcus 7GP-03_6His**

E. coli BL21 Star (DE3) competent cells were co-transformed with pACYC_ho1_pcyA (Set A) or pPcyA (Set B) and pET-R_7GP-03-6His. Recombinant proteins were then expressed and purified using $Ni^{2+}$ -affinity chromatography. A, B: SDS-PAGE gels of protein samples from the two independent transformants (Set A and Set B, respectively) taken at the various points during the purification process and stained with Coomassie blue. C: Separate SDS-PAGE gel of the elution fractions stained with Coomassie blue. D: Zinc fluorescence assay of the elution fractions. E: Overlay of a Western blot assay of the elution fractions. Western blotting was performed using antibodies specific for the 6xHis tag on the protein and visualized using HRP. F: Ponceau stain of the Western blot membrane shown in E. MW: molecular weight marker, Ho1, PcyA T0: transformed without pET-R_7GP-03-6His (before induction), Ho1, PcyA T1: transformed without pET-R_7GP-03-6His (induced for 16 hours), 7GP-03-6His T0: transformed only with pET-R_7GP03-6His (before induction), 7GP-03_6His T1: transformed only with pET-R_7GP-03-6His (induced for 16 hours), Both T0: transformed with both plasmids from the set (before induction), Both T1: transformed with both plasmids from the set (induced for 16 hours), Load: supernatant from cell lysate (before adding to affinity column), FT: supernatant from cell lysate (after flowing through the affinity column), W: column flowthrough after applying wash buffer, E1-E5: eluates in 2 mL increments after applying the elution buffer. For further details see Methods.

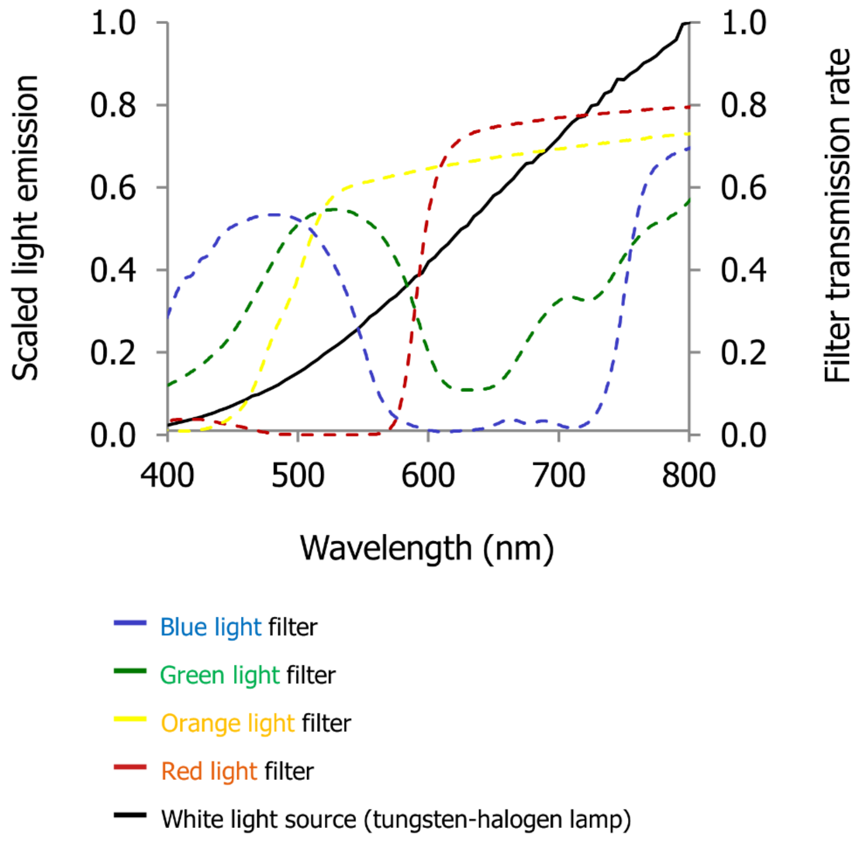


SUPPL. FIG. 2: **Spectra of colour treatments**

The tansmission (%T) spectra of four colour filters were recorded using an absorbance spectrometer. Lines are coloured and labelled according to the largest wavelength band the filter transmits, namely blue [370:560 nm], green [420:620 nm], orange [430:∞), red [570:∞). The emission spectrum of the tungsten-halogen white light lamp is also measured and shown in black.

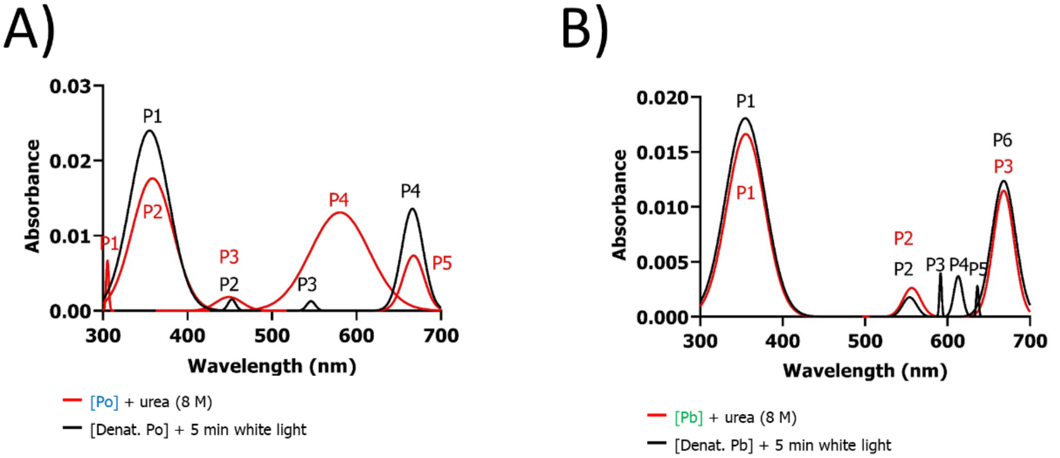


C)

| | Peak | Location (nm) | Height | Width | Area |
|---|---|---|---|---|---|
| Denat. [Po] | P1 | 305.38 | 0.01 | 2.2 | 0.03 |
| | P2 | 358.69 | 0.02 | 34.18 | 1.07 |
| | P3 | 449.11 | 0 | 21.65 | 0.07 |
| | P4 | 580.41 | 0.01 | 49.17 | 1.14 |
| | P5 | 667.61 | 0.01 | 17.58 | 0.23 |
| Denat. [Po] + white light | P1 | 355.2 | 0.02 | 33.16 | 1.41 |
| | P2 | 452.47 | 0 | 6.54 | 0.02 |
| | P3 | 546.18 | 0 | 7.25 | 0.02 |
| | P4 | 666.22 | 0.01 | 17.99 | 0.43 |

D)

| | Peak | Location (nm) | Height | Width | Area |
|---|---|---|---|---|---|
| Denat. [Pb] | P1 | 355.26 | 0.02 | 32.62 | 0.96 |
| | P2 | 556.6 | 0 | 15.02 | 0.07 |
| | P3 | 668.07 | 0.01 | 17.75 | 0.36 |
| Denat. [Pb] + white light | P1 | 354.61 | 0.02 | 33.87 | 1.09 |
| | P2 | 553.98 | 0 | 12.13 | 0.04 |
| | P3 | 591.5 | 0 | 1.93 | 0.01 |
| | P4 | 612.87 | 0 | 7.28 | 0.05 |
| | P5 | 696.27 | 0 | 2.06 | 0.01 |
| | P6 | 668.08 | 0.01 | 20.36 | 0.45 |

SUPPL. FIG. 3: **Peak deconstruction for Figure 3**

Recombinant 7GP-03 was expressed in the presence of PCB and purified as a blue solution under standard light conditions. Aliquots of the solution were exposed for 5 minutes to 'blue' or 'orange' light to convert all protein to Po (blue line) or Pb (green line), respectively. Urea was then added to the samples, followed by white light illumination for 5 min. A and B: Peak deconstruction of the absorbance spectra from the denatured samples in Figure 4a and b, respectively. The spectra taken immediately after urea denaturation are shown in red, the spectra taken after white light illumination are shown in black. Peak deconstruction was performed as described in the Methods.

A)

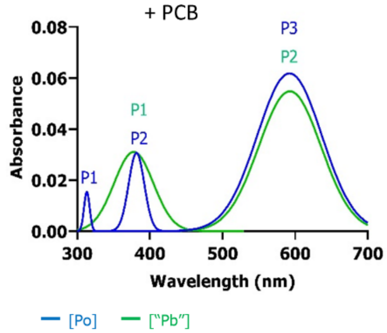


B)

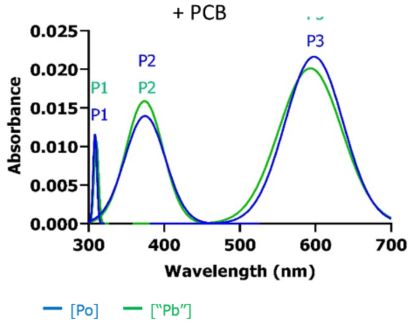


C)

| | Peak | Location (nm) | Height | Width | Area |
|---|---|---|---|---|---|
| [Po] | P1 | 313.29 | 0.02 | 5.43 | 0.15 |
| | P2 | 381.62 | 0.03 | 15.2 | 0.82 |
| | P3 | 592.31 | 0.06 | 62.86 | 6.89 |
| ["Pb"] | P1 | 377.82 | 0.03 | 38.17 | 2.1 |
| | P2 | 592.83 | 0.05 | 60.59 | 5.88 |

D)

| | Peak | Location (nm) | Height | Width | Area |
|---|---|---|---|---|---|
| [Po] | P1 | 308.5 | 0.01 | 3.47 | 5.78 |
| | P2 | 374.61 | 0.01 | 38.02 | 63.31 |
| | P3 | 598.22 | 0.02 | 54.37 | 90.54 |
| ["Pb"] | P1 | 310.15 | 0.01 | 4.14 | 0.08 |
| | P2 | 373.9 | 0.02 | 34.87 | 0.98 |
| | P3 | 593.33 | 0.02 | 58.87 | 2.1 |

SUPPL. FIG. 4: Peak deconstruction from Figure 4

Mutant versions of 7GP-03 replacing cysteine in position 73 with alanine (7GP-03_C73A) or serine (7GP-03_C73S) were generated by site-directed mutagenesis, the recombinant proteins were expressed In the presence or absence of PCB and purified under standard light conditions. Absorbance spectroscopy was then carried out after treating the protein with 'blue' light (blue lines) or 'orange' light for 1 minute or 5 minutes. A, B: The results of peak deconstruction of the 7GP-03_C73A and 7GP-03_C73S spectra from Figure 5. C, D: the data from A) and B), respectively.

A)

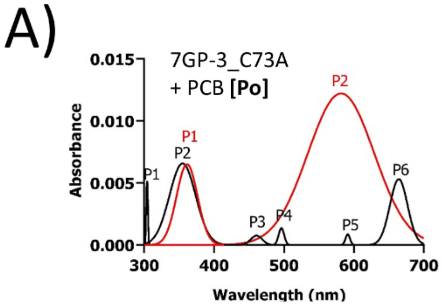


B)

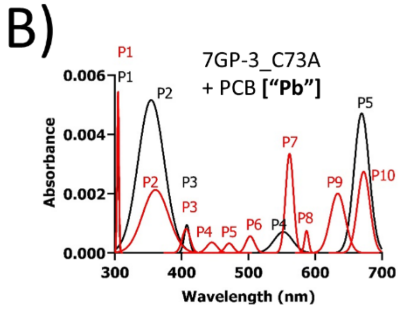


C)

Denatured — Denatured + white light

| | Peak | Location (nm) | Height | Width | Area |
|---|---|---|---|---|---|
| Den. [Po] | P1 | 361.37 | 0.01 | 21.21 | 0.24 |
| | P2 | 581.85 | 0.01 | 67.04 | 1.45 |
| Den. [Po] + white light | P1 | 304.12 | 0.01 | 1.68 | 0.02 |
| | P2 | 354.97 | 0.01 | 26.55 | 0.31 |
| | P3 | 460.87 | 0 | 11.59 | 0.02 |
| | P4 | 496.46 | 0 | 5.49 | 0.01 |
| | P5 | 591.42 | 0 | 4.54 | 0.01 |
| | P6 | 664.42 | 0.01 | 17.82 | 0.17 |

D)

Denatured — Denatured + white light

| | Peak | Location (nm) | Height | Width | Area |
|---|---|---|---|---|---|
| Den. [Pb] | P1 | 304.74 | 0.01 | 1.93 | 0.02 |
| | P2 | 360.83 | 0 | 27.5 | 0.1 |
| | P3 | 406.85 | 0 | 8.54 | 0.01 |
| | P4 | 445.29 | 0 | 10.92 | 0.01 |
| | P5 | 471.23 | 0 | 8.83 | 0.01 |
| | P6 | 502.69 | 0 | 9.23 | 0.01 |
| | P7 | 561.67 | 0 | 8.29 | 0.05 |
| | P8 | 587.19 | 0 | 4.31 | 0.01 |
| | P9 | 633.9 | 0 | 15.75 | 0.06 |
| | P10 | 672.73 | 0 | 13.09 | 0.06 |
| Den. [Pb] + white light | P1 | 304.58 | 0 | 1.87 | 0.02 |
| | P2 | 354.32 | 0.01 | 27.56 | 0.25 |
| | P3 | 407.79 | 0 | 5.62 | 0.01 |
| | P4 | 552.26 | 0 | 19.62 | 0.02 |
| | P5 | 669.76 | 0 | 15.96 | 0.13 |

E)

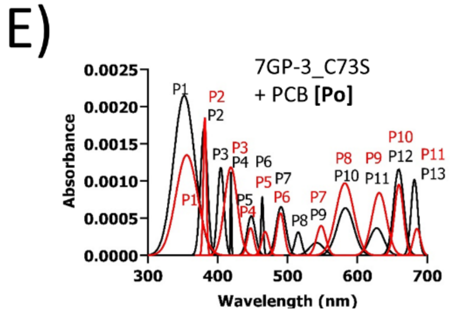


F)

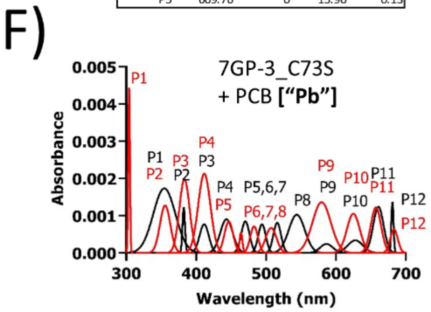


Denatured — Denatured + white light

Denatured — Denatured + white light

G)

| | Peak | Location (nm) | Height | Width | Area |
|---|---|---|---|---|---|
| Den. [Po] | P1 | 355.71 | 0 | 21.842 | 0.05 |
| | P2 | 381.64 | 0 | 3.01 | 0.01 |
| | P3 | 418.95 | 0 | 12.92 | 0.03 |
| | P4 | 447.2 | 0 | 6.8 | 0 |
| | P5 | 467.93 | 0 | 7 | 0 |
| | P6 | 490 | 0 | 7.37 | 0.01 |
| | P7 | 548.18 | 0 | 9.33 | 0.01 |
| | P8 | 582.16 | 0 | 18.09 | 0.03 |
| | P9 | 631.32 | 0 | 14.83 | 0.2 |
| | P10 | 659.56 | 0 | 9.03 | 0.02 |
| | P11 | 685.34 | 0 | 7.32 | 0 |
| Den. [Po] + white light | P1 | 352.12 | 0 | 21.36 | 0.08 |
| | P2 | 380.46 | 0 | 7.07 | 0.02 |
| | P3 | 404.44 | 0 | 6.47 | 0.01 |
| | P4 | 419.04 | 0 | 1.06 | 0 |
| | P5 | 448.07 | 0 | 8.34 | 0.01 |
| | P6 | 463.83 | 0 | 2.35 | 0 |
| | P7 | 490.7 | 0 | 9.67 | 0.01 |
| | P8 | 515.35 | 0 | 6.66 | 0 |
| | P9 | 541.58 | 0 | 14.16 | 0 |
| | P10 | 582.95 | 0 | 16.75 | 0.02 |
| | P11 | 627.77 | 0 | 14.23 | 0.01 |
| | P12 | 659.24 | 0 | 10.33 | 0.02 |
| | P13 | 681.74 | 0 | 6.79 | 0.01 |

H)

| | Peak | Location (nm) | Height | Width | Area |
|---|---|---|---|---|---|
| Den. [Pb] | P1 | 304.09 | 0 | 1.67 | 0.01 |
| | P2 | 355.65 | 0 | 10.43 | 0.02 |
| | P3 | 383.31 | 0 | 10.31 | 0.04 |
| | P4 | 411.66 | 0 | 12.32 | 0.05 |
| | P5 | 446.68 | 0 | 8.7 | 0.01 |
| | P6 | 464.15 | 0 | 2.94 | 0 |
| | P7 | 482.88 | 0 | 7.76 | 0.01 |
| | P8 | 507.23 | 0 | 10.92 | 0.01 |
| | P9 | 579.55 | 0 | 18.5 | 0.04 |
| | P10 | 624.93 | 0 | 13.04 | 0.02 |
| | P11 | 657.2 | 0 | 10.4 | 0.02 |
| | P12 | 683.09 | 0 | 7.65 | 0.01 |
| Den. [Pb] + white light | P1 | 354.52 | 0 | 22.33 | 0.07 |
| | P2 | 382.28 | 0 | 2.56 | 0.01 |
| | P3 | 411.47 | 0 | 9.3 | 0.01 |
| | P4 | 443.43 | 0 | 12.59 | 0.02 |
| | P5 | 470.43 | 0 | 8.3 | 0.01 |
| | P6 | 494.24 | 0 | 7.58 | 0.01 |
| | P7 | 515.99 | 0 | 7.72 | 0.01 |
| | P8 | 543.98 | 0 | 16.11 | 0.03 |
| | P9 | 586.49 | 0 | 13 | 0.01 |
| | P10 | 627.8 | 0 | 15.32 | 0.01 |
| | P11 | 660.75 | 0 | 10.48 | 0.02 |
| | P12 | 680.86 | 0 | 2.24 | 0.01 |

SUPPL. FIG. 5: Peak deconstruction from Figure 5

Mutant versions of 7GP-03 replacing cysteine in position 73 with alanine (7GP-03_C73A) or serine (7GP-03_C73S) were generated by site-directed mutagenesis, the recombinant proteins were expressed in the presence or absence of PCB and purified under standard light conditions. Aliquots of the solution were exposed for 5 minutes to 'blue' or 'orange' light to try and convert all protein to Po or Pb, respectively. A, B: Absorbance spectra deconstruction of 7GP-03_C73A in "Po" state (A) or "Pb" state (B) after addition of 8 M urea (red) and immediately after white light illumination (black). C, D: the peak deconstruction data from A) and B), respectively. E, F: Absorbance spectra deconstruction of 7GP-03_C73A in "Po" state (A) or "Pb" state (B) after addition of 8 M urea (red) and immediately after white light illumination (black). G, H: the peak deconstruction data from E) and F), respectively.

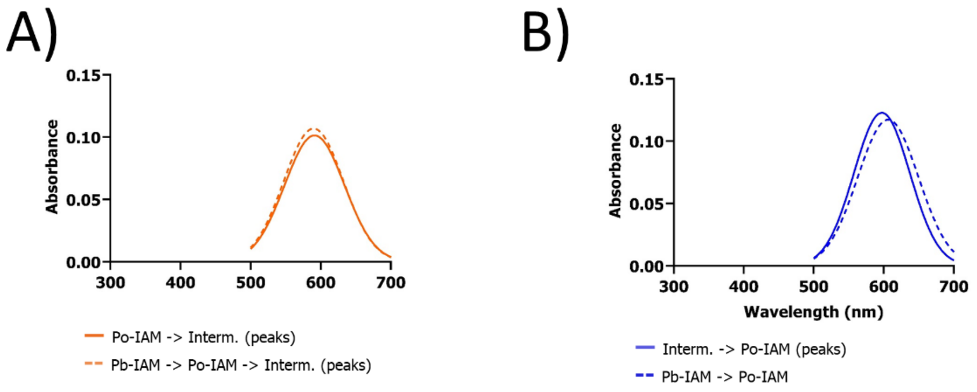


C)

| | Peak | Location (nm) | Height | Width | Area |
|---|---|---|---|---|---|
| Po-IAM -> Interm. | P1 | 592.27 | 0.1 | 60.5 | 10.88 |
| Pb-IAM -> Po-IAM -> Interm. | P1 | 589.96 | 0.11 | 60.19 | 11.41 |

D)

| | Peak | Location (nm) | Height | Width | Area |
|---|---|---|---|---|---|
| Interm. -> Po-IAM | P1 | 597.44 | 0.12 | 56.55 | 12.31 |
| Pb-IAM -> Po-IAM | P1 | 606.36 | 0.12 | 61.43 | 12.77 |

SUPPL. FIG. 6: **Peak deconstruction from Figure 6**

Recombinant 7GP-03 was expressed in the presence of PCB and purified as a blue solution under standard light conditions. Aliquots of the solution were exposed for 5 minutes to 'blue' or 'yellow' light to convert all protein to Po or Pb, respectively. IAM was then added to a final concentration of 50 mM. Next, the samples were exposed to 5 min of 'orange' or 'blue' light to drive the protein into the opposite state. Finally, the protein was motivated back to its original state by applying either 'blue' or 'orange' light. A: the deconstructed peaks of the IAM-blocked Po -> Pb conversion (Interm.). The continuous line shows the Po -> Pb direction. The dashed line shows the Pb -> Po -> Pb direction. B: The deconstructed peaks of the opposite direction are shown. The continuous line shows the Interm. -> Po-IAM direction. The dashed line shows the Pb-IAM -> Po-IAM direction. C, D: the data of the deconstructed peaks from A) and B), respectively.

A)

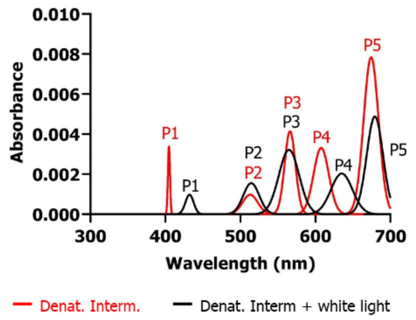


B)

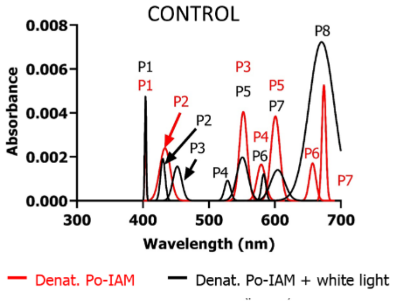


C)

| | Peak | Location (nm) | Height | Width | Area |
|---|---|---|---|---|---|
| Denat. Interm. | P1 | 404.78 | 0 | 1.95 | 0.01 |
| | P2 | 513.07 | 0 | 14.29 | 0.02 |
| | P3 | 566.17 | 0 | 10.164 | 0.07 |
| | P4 | 607.75 | 0 | 14.35 | 0.08 |
| | P5 | 674.32 | 0.01 | 14.08 | 0.2 |
| Denat. Interm. + white light | P1 | 432.17 | 0 | 7.74 | 0.01 |
| | P2 | 514.49 | 0 | 15.02 | 0.04 |
| | P3 | 564.81 | 0 | 19.15 | 0.11 |
| | P4 | 635.11 | 0 | 20.23 | 0.07 |
| | P5 | 679.3 | 0 | 15.41 | 0.13 |

D)

| | Peak | Location (nm) | Height | Width | Area |
|---|---|---|---|---|---|
| Denat. Po-IAM | P1 | 403.89 | 0 | 1.59 | 0.01 |
| | P2 | 433.48 | 0 | 11.28 | 0.05 |
| | P3 | 552.15 | 0 | 8.44 | 0.06 |
| | P4 | 579.33 | 0 | 9.1 | 0.03 |
| | P5 | 600.82 | 0 | 9.57 | 0.07 |
| | P6 | 657.2 | 0 | 6.98 | 0.07 |
| | P7 | 674.32 | 0 | 4.25 | 0.02 |
| Denat. Po-IAM + white light | P1 | 403.93 | 0 | 1.6 | 0.01 |
| | P2 | 430.18 | 0 | 4.98 | 0.02 |
| | P3 | 452.07 | 0 | 9.29 | 0.03 |
| | P4 | 528.28 | 0 | 6.52 | 0.01 |
| | P5 | 550.78 | 0 | 11.86 | 0.04 |
| | P6 | 583.02 | 0 | 4.73 | 0.01 |
| | P7 | 604.34 | 0 | 14.94 | 0.04 |
| | P8 | 670.85 | 0 | 27.05 | 0.35 |

SUPPL. FIG. 7: **Peak deconstruction from Figure 8**

Recombinant 7GP-03 was expressed in the presence of PCB and purified as a blue solution under standard light conditions. An aliquot of the protein was exposed to 5 min 'blue' light to take it completely to Po. IAM was then added to a final concentration of 50 mM. The IAM- treated sample was then illuminated for 5 min with 'orange' light to push the protein in the Po -> Pb direction. Next, urea was added to a final concentration of 8 M, followed by white light treatment for 5 min. A: The deconstructed peaks from Figure 8, where red indicates the peaks immediately after urea-based denaturation and black indicates the peaks after white light illumination. C, D: the data of the deconstructed peaks from A) and B), respectively.

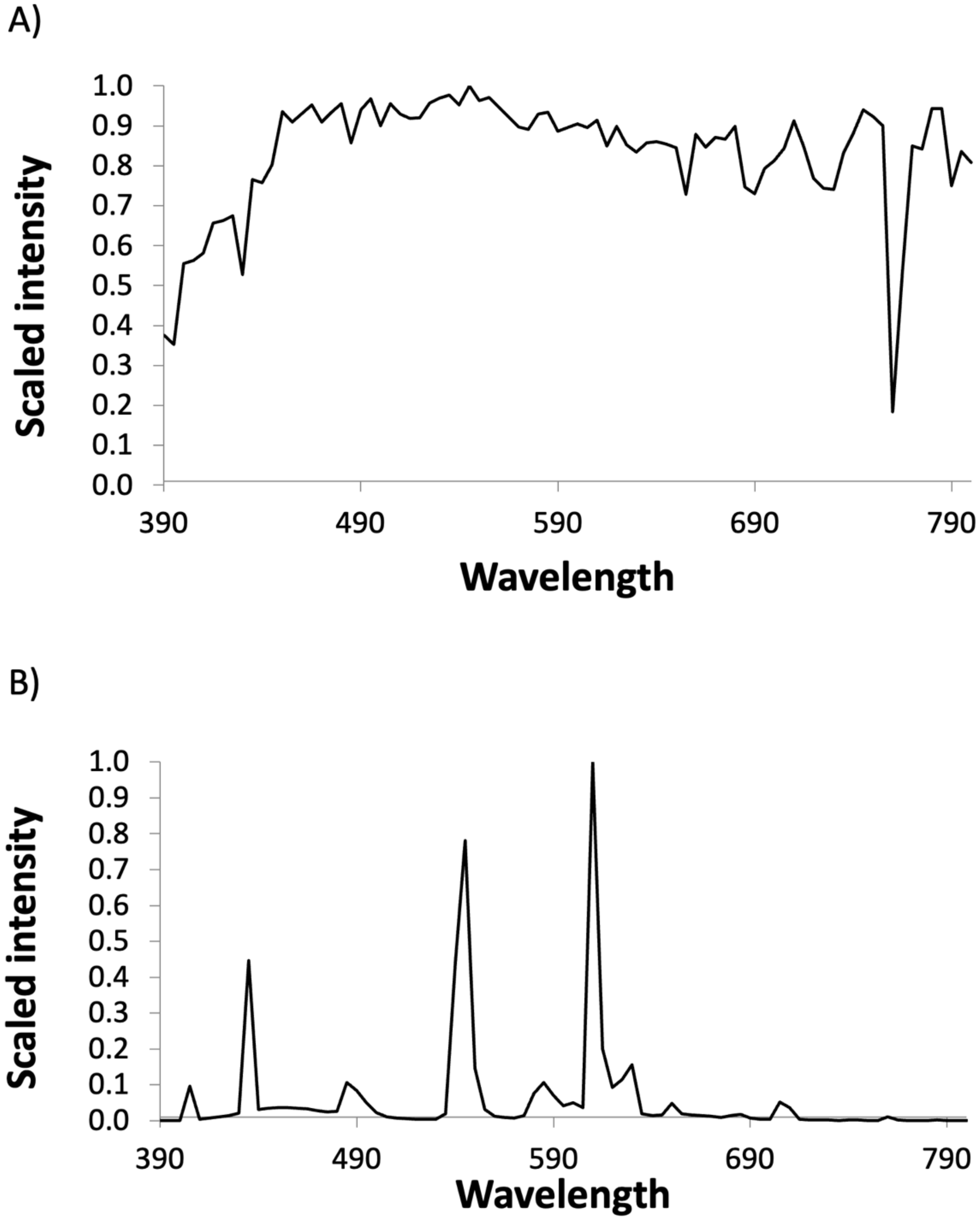


SUPPL. FIG. 8: **Emission spectra mixed into the ambient light environment**

The emission spectra were measured separately using a directional quartz/optic fiber probe, pointed at the light source. Measurements were done using a spectroradiometer (unknown brand). Power output was measured in µE and subsequently scaled and shown. A: Power output from the sun on a typical day (cloudy), across the visible spectrum. Measured through a window at noon with all other lights off. B: Power output from the warm-white triphosphor fluorescent tubes, used to illuminate the lab (measured at night).
